# Whole-brain estimates of directed connectivity for human connectomics

**DOI:** 10.1101/2020.02.20.958124

**Authors:** Stefan Frässle, Zina M. Manjaly, Cao T. Do, Lars Kasper, Klaas P. Pruessmann, Klaas E. Stephan

## Abstract

Connectomics is essential for understanding large-scale brain networks but requires that individual connection estimates are neurobiologically interpretable. In particular, a principle of brain organization is that reciprocal connections between cortical areas are functionally asymmetric. This is a challenge for fMRI-based connectomics in humans where only undirected functional connectivity estimates are routinely available. By contrast, whole-brain estimates of effective (directed) connectivity are computationally challenging, and emerging methods require empirical validation.

Here, using a motor task at 7T, we demonstrate that a novel generative model can infer known connectivity features in a whole-brain network (>200 regions, >40,000 connections) highly efficiently. Furthermore, graph-theoretical analyses of directed connectivity estimates identify functional roles of motor areas more accurately than undirected functional connectivity estimates. These results, which can be achieved in an entirely unsupervised manner, demonstrate the feasibility of inferring directed connections in whole-brain networks and open new avenues for human connectomics.

## 1 INTRODUCTION

Understanding the human brain is a major scientific challenge of our time. Recent methodological advances have provided unprecedented opportunities for studying the brain^1–3^. In particular, non-invasive neuroimaging techniques, such as functional magnetic resonance imaging (fMRI), have enabled studying the living human brain as a dynamic system of interconnected neuronal populations^4^. This has fueled the emergence of whole-brain connectomics, a young discipline which is fundamentally important for understanding the organizational principles of the brain and plays a central role in network neuroscience^5^.

Since the term “connectome” was originally introduced^6, 7^, the field has grown rapidly and is now one of the most vibrant disciplines in neuroscience^8^. One of the goals of connectomics is a comprehensive map of neuronal connections, covering the entire nervous system. Seminal achievements include the specification of the complete neuronal wiring diagram in *C. elegans*^9^ or the visual system of *Drosophila*^10^. In non-human primates and humans, particular emphasis has been placed on differences and individuality. For example, an important concept is that of “connectivity fingerprints” – a term originally introduced to refer to area-specific patterns of connectivity^11^ and more recently used to denote subject-specific connectivity patterns that determine inter-individual differences in brain function^12^ and behavior^13^. Furthermore, connectomics has begun incorporating changes in connectivity with cognitive context or learning^14^.

Connectomics is not only crucial for studying organizational principles in the healthy human brain, but also in disease. Aberrant functional integration has been observed in most psychiatric and neurological disorders^15–18^. For example, psychiatric diseases like schizophrenia, depression, and autism have all been associated with pathological alterations across the functional connectome. For this reason, connectomes may serve as intermediate phenotypes situated between the domains of genetics/molecules and expressions of individual (pathological) behavior^15^.

However, to render connectomics useful for understanding large-scale brain networks and alterations thereof, individual connection estimates have to be neurobiologically interpretable. A principle of brain organization are functional asymmetries of reciprocal connections – for instance, differences between ascending and descending connections in cortical hierarchies^19, 20^ or asymmetries in interhemispheric interactions^21–23^. This however represents a challenge for fMRI-based connectomics in humans: routine measures of connectivity are so far undirected; namely, structural and functional connectivity among network nodes at a mesoscopic or macroscopic level. In brief, structural connectivity refers to white-matter fiber tracts that can be measured using diffusion weighted imaging (DWI)^24^, whereas functional connectivity relates to statistical interdependencies between fMRI signals and is computed using simple correlation analyses or more sophisticated statistical techniques (for a comprehensive review, see ^25^).

Unfortunately, inferring directed estimates of functional interactions (i.e., effective connectivity) at the whole-brain level has proven challenging, mainly due to computational limitations. While various models of effective connectivity have been proposed over the last decade^26^, including dynamic causal models (DCMs)^27^ and biophysical network models (BNMs)^28, 29^, these methods are limited in either the network size that can be considered (DCM) or the ability to identify individual connection strengths (BNM). While recent progress has been made in both domains^30–32^, computational efficiency and identifiability remain problematic and/or unknown. In addition to methodological refinements, basic empirical validation studies are required that challenge models to rediscover known sets of connections from whole-brain fMRI data.

We have recently introduced a novel method that enables connection-specific estimates of effective connectivity in whole-brain networks. This method – termed regression dynamic causal modeling (rDCM)^33, 34^ – is promising for several reasons: First, rDCM is computationally highly efficient and scales gracefully to large networks that comprise hundreds of nodes. Second, the model can exploit structural connectivity information to constrain inference on directed functional interactions or, where no such information is available, infer optimally sparse representations of whole-brain networks. Hence, rDCM provides two alternative modes of operation to derive individual connectivity fingerprints at the whole-brain level. Third, rDCM allows to exploit knowledge about where and when experimental perturbations (e.g., sensory stimuli) affect network dynamics. This is important since known perturbations can greatly help constrain inference about directed influences within systems^35^.

In this paper, we illustrate the practical benefits of rDCM for whole-brain connectomics and network neuroscience in humans. For this, we use ultra-high field 7T fMRI data acquired under a deliberately simple paradigm (visually paced hand movements) in which relevant connections are well known and show clear hemispheric asymmetries. Lateralized processes are particularly useful for this purpose as they provide strong qualitative predictions^21, 23^ that concern both the location (hemisphere) where processes should occur (or, equally important, not occur) as well as the asymmetry (or mirror symmetry) of processes across hemispheres. Here, we demonstrate the utility of rDCM by performing two types of whole-brain connectivity analyses in a network with over 200 regions and 40,000 directed connections. These analyses are (i) anatomically guided by tractography results, and (ii) completely unconstrained by pruning fully (all-to-all) connected brain-wide graphs to those connections essential for explaining whole-brain activity.

## 2 METHODS AND MATERIALS

### 2.1 Regression dynamic causal modeling

#### 2.1.1 Basic framework

Regression DCM (rDCM) is a novel variant of DCM for fMRI that has specifically been developed for effective connectivity analyses in large (whole-brain) networks^34^. For this, rDCM applies several modifications and simplifications to the original DCM framework (for a short summary of classical DCM, see Supplementary Material S1). In brief, these include (i) translating state and observation equations from time to frequency domain using the Fourier transformation (under stationarity assumptions), (ii) replacing the nonlinear biophysical model of hemodynamics with a linear hemodynamic response function (HRF), (iii) applying a mean field approximation across regions (i.e., connectivity parameters targeting different regions are assumed to be independent), and (iv) specifying conjugate priors on neuronal (i.e., connectivity and driving input) parameters and noise precision to enable analytic variational Bayesian (VB) update equations. These modifications essentially transform a linear DCM in the time domain into a Bayesian linear regression in the frequency domain:

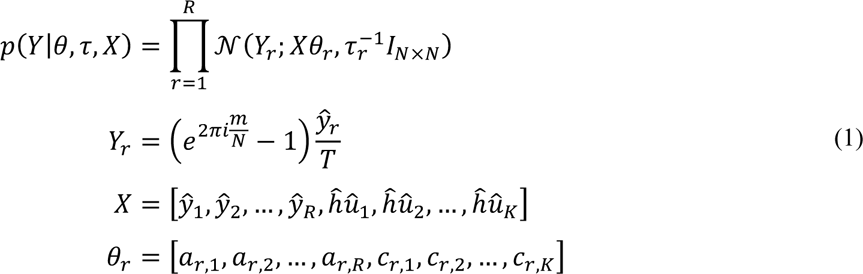

where *Y*_*r*_ is the signal in region *r* that is explained as a linear mixture of afferent connections from other regions and direct (driving) inputs, *y*_*r*_ is the measured BOLD signal in region *r*, *X* is the design matrix (comprising a set of regressors and explanatory variables), and *u*_k_ is the k^th^ experimental input. Furthermore, *λ*_*r*_ represents the parameter vector comprising all connections *a*_*r,1*_, … *a*_*r,R*_ and all driving input parameters *c*_*r,*10_, …, *c*_*r,K*_ targeting region *r*. Finally, τ_*r*_ denotes the noise precision parameter for region *r* and *I*_*N×N*_ is the identity matrix (where *N* denotes the number of data points). Under this formulation, inference can be done very efficiently by (iteratively) executing a set of analytical VB update equations concerning the sufficient statistics of the posterior density. In addition, one can derive an expression for the negative (variational) free energy^36^. The negative free energy represents a lower-bound approximation to the log model evidence that accounts for both model accuracy and complexity. Hence, the negative free energy offers a sensible metric for scoring model goodness and thus serves as a criterion for comparing competing hypotheses^37^. A comprehensive description of the generative model underlying rDCM can be found elsewhere^34^.

#### 2.1.2 Sparsity constraints

The standard rDCM framework has recently been augmented with sparsity constraints to enable automatic pruning of fully (all-to-all) connected networks to a degree of optimal sparsity^33^. This is achieved by introducing an additional set of binary indicator variables as feature selectors in the likelihood function. In particular, each connectivity and driving input parameter *i* in a fully connected model is multiplied with a specific binary indicator variable ξ_*i*_ which takes the value of 1 if the connection is present (i.e., contributes to explaining the observed signal) and 0 if the connection is absent (i.e., not involved in generating the observed signal). The Bayesian sparse linear regression model in the frequency domain takes the form:

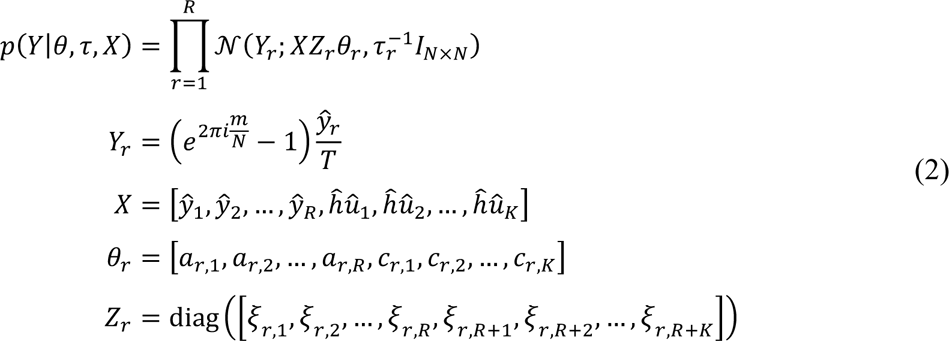

Where *Z*_*r*_ is a diagonal matrix comprising the binary indicator variables for all connections and driving inputs targeting region *r*. All other variables in Eq. (2) are defined as above. For this framework, one can again derive a VB update scheme for model inversion to obtain estimates of (i) the posterior distribution over neuronal connectivity, noise precision and binary indicator parameters, and (ii) the negative free energy. As in the basic rDCM framework, model inversion then boils down to iteratively solving a set of update equations. For a comprehensive description of the generative model, we refer the reader to previous work^33^.

### 2.2 Empirical data

Here, we assess the utility of rDCM for inferring the effective connectivity pattern in a realistic whole-brain network based on empirical data from an fMRI study with a simple paradigm of visually paced hand movements. We chose this dataset for the following reasons: (i) the simple and robust nature of the task, (ii) the extensive knowledge available about the cortical network supporting hand movements^38–40^, (iii) the engagement of distributed cortical networks related to visual and motor aspects of the task, (iv) the high signal-to-noise ratio (SNR) of the data afforded by the 7 Tesla MR scanner on which the data was acquired, and (v) the absence of modulatory influences, which appeals to the linearity assumptions in rDCM. This allowed us to probe the plausibility of the whole-brain connectivity patterns inferred by rDCM.

#### 2.2.1 Participants

Thirty right-handed individuals (14 females, 16 males; mean age: 59.2 ± 9.5 years, age range: 39-74 years) participated in this methodological study. Five participants had to be excluded from the analysis due to non-compliance with the task, missing data, or incorrect scanner settings, resulting in a final sample of 25 participants (13 females, 12 males; mean age: 58.2 ± 9.7 years, age range: 39-71 years). All were healthy with no history of psychiatric or neurological disease, brain pathology or abnormalities in brain morphology as indicated by their T1-weighted anatomical image. All participants were fluent German speakers. Half of the participants were regularly taking low-dose aspirin (100mg per day). This is because the dataset used in this paper consists of two groups from a larger study that was purely observational (i.e., participants already took aspirin independently from our study). In the following, we treat participants with and without aspirin intake as one group, given that the present study is not interested in potential aspirin effects but merely serves to test the construct validity of rDCM for a simple paradigm. For completeness, we examined potentially confounding effects of aspirin on connectivity estimates and found that none of the whole-brain connectivity estimates presented below showed any significant differences between the two groups. For each participant, written informed consent was obtained prior to the experiment. The experimental protocol was in compliance with the Declaration of Helsinki and was performed with approval by the Cantonal Ethics Commission of Zurich (EK 09-2006 ETH).

#### 2.2.2 Experimental procedure

The task used in this study was similar to the paradigm by Grefkes et al.^41^ in which participants had to perform visually synchronized whole-hand fist closings with either their left or right hand. Hand movements of different conditions (i.e., left or right) were separated into two scanning sessions (which deviates from the work by Grefkes and colleagues, where left- and right-hand movements were performed in alteration within a single session).

In each session, the experimental paradigm was a blocked design as follows: At the beginning of each block, an arrow was presented in the middle of the screen, indicating which hand to use in the upcoming block. The arrow then started blinking at a rate of 1.25Hz for 16s, dictating the rhythm of participants’ hand movements (i.e., 20 fist closings per block). The stimulus onset time was 300ms and the inter-stimulus interval was set to 500ms. Subsequent blocks were interleaved with a resting period of the same length where participants did not perform hand movements but kept fixation in the center of the screen. Stimuli were presented using Cogent 2000 (v1.33; http://www.vislab.ucl.ac.uk/Cogent/index.html). Since each session comprised fist closings of only one condition, this dataset is particularly suitable for probing the current implementation of rDCM since no modulatory influences are required.

As mentioned before, the task was chosen because it affords clear hypotheses about the putative network supporting visually synchronized hand movements^38–40^. Specifically, simple unilateral hand movements (i) result from lateralized brain activity^42^ and (ii) involve interactions between well-known brain regions. We briefly comment on these properties in more detail.

Lateralized processes are particularly useful to evaluate models of connectivity as they provide strong qualitative predictions (for previous examples, see ^21, 23^). These predictions concern the location (hemisphere) where processes should occur (or, equally important, not occur) as well as the asymmetry (or mirror symmetry) of processes across hemispheres. In our paradigm, unilateral hand movements should be accompanied by enhanced connectivity between motor areas in the contralateral hemisphere^43^. Given the visual pacing input, one would also expect lateralized connectivity from visual (e.g., motion-sensitive area V5/MT) to motor areas (via parietal areas)^38, 39^; the effect of lateralization may be less strong, however, since the visual input was presented centrally and thus did not specifically enter one hemisphere. In the motor domain, an additional advantage of our paradigm is that contrasting left and right unilateral hand movements allows for mirror-symmetric predictions: right-hand movements should lead to enhanced connectivity between left-hemispheric (but not the corresponding right-hemispheric) motor regions and vice versa. This offers an opportunity to test the replicability of our connectivity findings across hemispheres.

The key cortical components of the motor network underlying visually synchronized unilateral hand movements are well known^40, 41^. These include primary motor (M1) and somatosensory cortex (SM1), supplementary motor area (SMA), and lateral premotor cortex (PMC). In brief, M1 represents the main executive locus, with corticospinal projections which directly target lateral motor nuclei in the spinal cord^44^. PMC is involved in the execution of hand movements under sensory guidance^45^, and was found to be crucial for transforming sensory information into appropriate motor behavior^46^. SMA represents an integral component for planning and initiating voluntary hand movements^41, 47^. Furthermore, SM1 relates to somatosensory and proprioceptive aspects of motor acts^48^. In addition to the components mentioned above, the anterior cerebellum is involved in simple unilateral hand movements^40^. Furthermore, given the visual pacing input, one would also expect visual areas such as the primary visual cortex and the motion-sensitive area V5/MT to be engaged.

#### 2.2.3 Data acquisition

Functional images were acquired using a 7T MR scanner (Philips Achieva) with a 16-channel head matrix receiver coil. Images were obtained using a T2*-weighted gradient echo-planar imaging (EPI) sequence (36 axial slices, TR=2000ms, TE=25 ms, field of view (FoV) 220×220×108mm^3^, voxel size 1.77×1.77×3mm^3^, flip angle 70°, SENSE factor 4) sensitive to the blood oxygen level dependent (BOLD) signal. Images covered the entire brain. In each session, 230 functional images were acquired, representing either brain activity during left- or right-hand fist closings. For each participant, an additional high-resolution anatomical image was acquired using a T1-weighted inversion-recovery turbo field echo (3D IR-TFE) sequence (150 slices, TR=7.7ms, TE=3.5ms, volume TR=4000ms, inversion time 1200ms, field of view (FoV) 240×240×135mm^3^, voxel size 0.9×0.9×0.9mm^3^, flip angle 7°, SENSE factor 2 in phase and 1.5 in slice direction).

In addition to the MRI data, physiological recordings related to heart beats and breathing were recorded during scanning with a 4-electrode electrocardiogram (ECG) and a breathing belt, respectively.

#### 2.2.4 Data processing and analysis

Functional images were analyzed using SPM12 (Statistical Parametric Mapping, version R6553, Wellcome Trust Centre for Neuroimaging, London, UK, http://www.fil.ion.ucl.ac.uk) and Matlab R2015a (Mathworks, Natick, MA, USA). Individual images were realigned to the mean image, unwarped, coregistered to the participant’s high-resolution anatomical image, and normalized to the Montreal Neurological Institute (MNI) standard space using the unified segmentation-normalization approach. During spatial normalization, functional images were resampled to a voxel size of 2×2×2mm^3^. Finally, normalized functional images were spatially smoothed using an 8mm FWHM Gaussian kernel.

Model-based physiological noise correction based on peripheral recordings of cardiac (heart beat) and respiratory (breathing) cycles was performed using the PhysIO toolbox^49^. Specifically, the periodic effects of pulsatile motion and field fluctuations were modeled using Fourier expansions (i.e., sine and cosine basis functions) of different order for the estimated phases (RETROICOR) of cardiac pulsation (3^rd^ order), respiration (4^th^ order) and cardio-respiratory interactions (1^st^ order). This resulted in 18 physiological noise regressors. The PhysIO toolbox is available as open source code as part of the TAPAS software suite (www.translationalneuromodeling.org/software).

Preprocessed functional images of every participant entered first-level General Linear Model analyses (GLM) to identify brain activity related to the experimental manipulation. The GLM comprised one task regressor, representing the periods when participants performed visually paced fist closings. The regressor was convolved with SPM’s standard canonical hemodynamic response function. Additionally, we included the temporal and dispersion derivative of the canonical HRF (“informed basis set”)^50^. To control for movement-related and physiological artefacts, respectively, motion parameters (obtained from rigid-body realignment of the functional volumes) and physiological measures (obtained from physiological noise modeling in PhysIO) were included as nuisance regressors in the GLM. Finally, a high-pass filter was applied to remove low-frequency fluctuations (e.g., scanner drifts) from the data (cut-off frequency: 1/128Hz).

Brain activity related to visually paced unilateral hand movements was then identified from the respective baseline contrasts of left- or right-hand movements. The individual contrast images were entered into random effects group level analyses (one-sample *t*-tests) for left- and right-hand fist closings, separately. Group-level BOLD activity was thresholded at *p*<0.05, family-wise error (FWE)-corrected at the peak level.

#### 2.2.5 Time series extraction

We used the Human Brainnetome atlas^51^ as a whole-brain parcellation scheme to define regions of interest for subsequent effective connectivity analyses. The Brainnetome atlas represents a connectivity-based parcellation derived from non-invasive structural neuroimaging data obtained from DWI (http://atlas.brainnetome.org). The atlas comprises 246 distinct parcels (123 per hemisphere), including 210 cortical and 36 subcortical regions. We chose the Brainnetome atlas as a parcellation scheme for the following reasons: (i) the atlas is sufficiently fine-grained to allow for meaningful effective connectivity analyses at the whole-brain level, (ii) provides robust parcels across the population as demonstrated using cross-validation, and (iii) includes not only a parcellation of the human brain but also information on the structural connectivity among the 246 brain regions. In a first analysis, we used this structural information to inform the architecture of our network – that is, the endogenous connectivity matrix. Notably, the Brainnetome atlas (like most other state-of-the-art parcellation schemes) focusses on the cortex and does not cover the cerebellum. Hence, in the present study, we made the deliberate choice to focus on the cortex in order to capitalize on the advantages of the Brainnetome atlas outlined above.

Due to signal dropouts in the raw functional images (especially in the pharahippocampal gyrus and inferior temporal regions near the skull base), BOLD signal time series could not be extracted for all regions defined by the Brainnetome atlas. In summary, 215 regions could be extracted in all participants for both hand movement conditions. We further restricted this set to ensure interhemispheric consistency of the network – that is, if a region was present in one hemisphere but not the other, both parcels were discarded from further analysis (for a complete list of included and excluded regions, see Supplementary Table S1). We verified that none of the excluded regions represented a key component of the cerebral network supporting visually paced hand movements^38–40^.

This yielded a total of 208 brain regions from which sensible BOLD signal time series could be obtained in every participant (for a visualization of the individual mean coordinates of each parcel, see Supplementary Figure S1). Time series were extracted as the principal eigenvariate of all voxels within a parcel. Time series were mean-centered and corrected for variance related to head movement, physiological noise, and derivatives of the hemodynamic response with regard to time and dispersion^50^. The latter served to address a limitation of the current rDCM implementation which employs a fixed hemodynamic response function and therefore does not capture hemodynamic variability across brain regions and individuals (see Discussion). Extracted BOLD signal time series then entered effective connectivity analysis using rDCM.

#### 2.2.6 rDCM analysis

For the rDCM analysis, we first used the structural connectome provided by the DWI data of the Brainnetome atlas to inform the connectivity architecture (i.e., the presence or absence of connections among brain regions in the A matrix) of the network (model 1; Figure 1A). As DWI data contains no information on the directionality of fibers, connected nodes were always coupled by reciprocal connections. Additionally, the driving input (representing visually synchronized left- or right-hand fist closing movements) was allowed to elicit activity in all regions. This yielded a total of 16,868 free parameters (including connectivity parameters, inhibitory self-connections and driving input parameters) to be estimated. To test the benefit of informing effective connectivity analyses by tractography-based measures, we further constructed two alternative networks: (i) a randomly permuted version of the Brainnetome structural connectome, discarding any regional specificity of connections while leaving the overall density of the network unchanged (model 2; Figure 1B), and (ii) a fully (all-to-all) connected network where all 208 brain regions are linked via reciprocal connections (model 3; Figure 1C).

**Figure 1:**
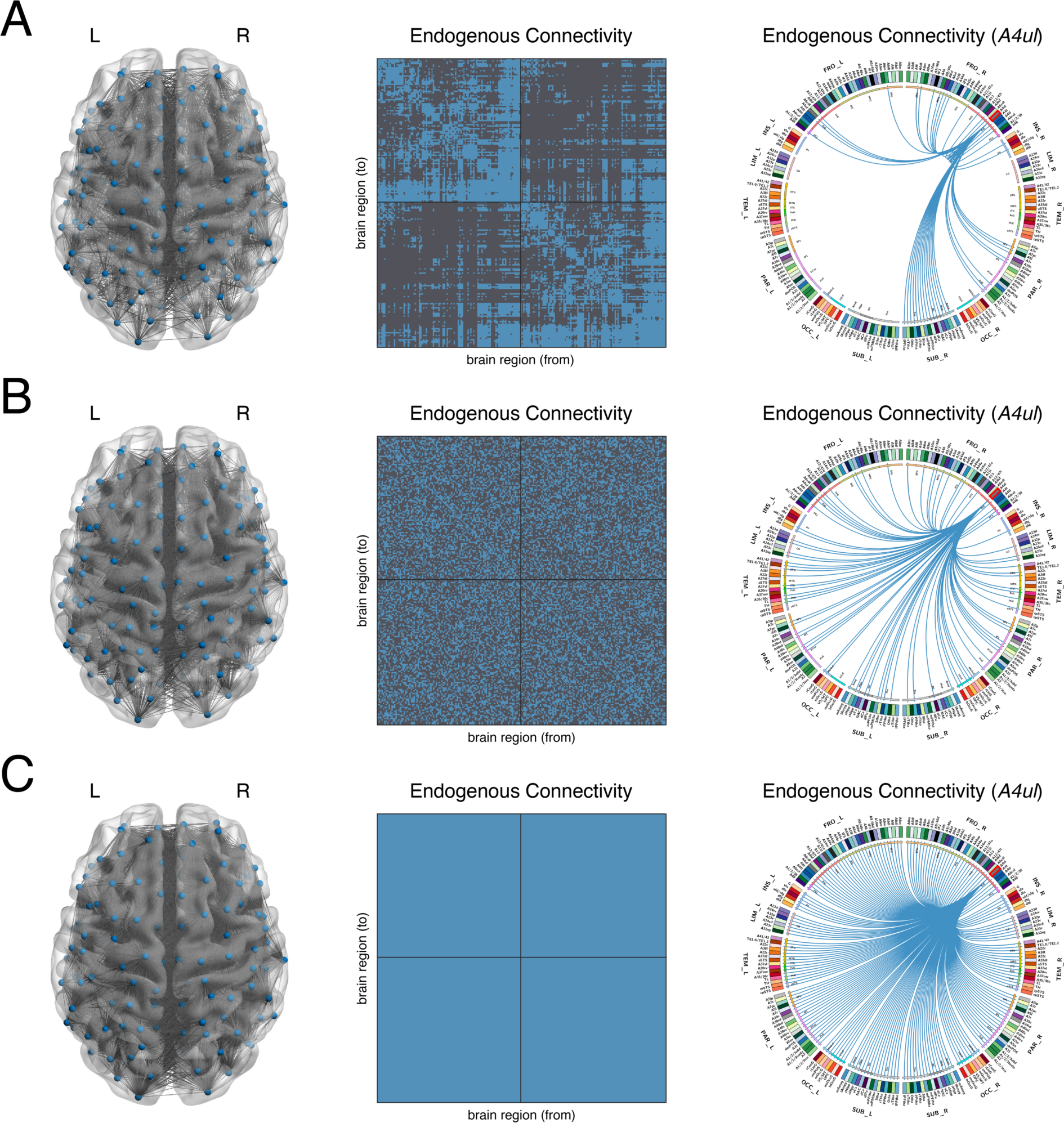
Connectivity architecture of the whole-brain networks used to model effective connectivity during unilateral hand movements. The alternative models encode a network architecture that **(A)** was informed by the structural connectome provided by the Human Brainnetome atlas (model 1), **(B)** was a randomly permuted version of the Brainnetome structural connectome; thus, discarding any regional specificity of connections while leaving the overall density of the network unchanged (model 2), or **(C)** was a fully (all-to-all) connected network, where all regions were reciprocally connected (model 3). For each of the three models, the network architecture of the DCM is graphically projected onto a whole-brain volume (*left*) and shown as an adjacency matrix (*middle*). Finally, we also show exemplarily the sub-regional connectogram for the primary motor cortex (M1) in the precentral gyrus (Brainnetome parcel name: *A4ul*) (*right*). The labels on the outermost ring of the connectogram show the anatomical lobe for each of the nodes: frontal, insula, limbic, temporal, parietal, occipital, and subcortical. For each brain region defined by the Brainnetome atlas, an abbreviation and color are defined. Inside the parcellation ring, we show the outgoing connections from M1 in blue. The whole-brain volume representation was created using the BrainNet Viewer^52^, which is freely available (http://www.nitrc.org/projects/bnv/). The connectogram was created using Circos, which is also publicly available (http://www.circos.ca/software/). L = left hemisphere; R = right hemisphere.

In a second step, we tested whether rDCM also yielded sensible results in the absence of any *a priori* restrictions on model architecture by utilizing the embedded sparsity constraints of the method to automatically prune both connections and driving inputs. To this end, we assumed a fully connected network, where all 208 brain regions were coupled to each other via reciprocal connections. Additionally, the driving input was again allowed to elicit activity in all regions. This yielded a total of 43,472 free parameters to be estimated. Starting from this fully connected network, model inversion then automatically pruned connection and driving input parameters to yield a sparse whole-brain effective connectivity pattern.

## 3 RESULTS

### 3.1 BOLD activity during unilateral hand movements

Brain activity related to visually synchronized whole-hand fist closings was assessed using random effects group analyses (one-sample *t*-tests). Consistent with previous findings, we observed significant activation in a widespread cortical network during left- and right-hand movements (Figure 2; Supplementary Table S2), mainly lateralized to the contralateral hemisphere. In particular, BOLD activation was located in the primary motor cortex (M1), premotor cortex (PMC), supplementary motor area (SMA), and the motion-sensitive area V5/MT in the extrastriate cortex (*p*<0.05, FWE-corrected at peak level). Additionally, we observed BOLD activation in the ipsilateral cerebellum. As mentioned before, for the subsequent effective connectivity analyses, we utilized the Brainnetome atlas^51^ as a whole-brain parcellation scheme which focuses on the cortex and does not cover the cerebellum.

**Figure 2:**
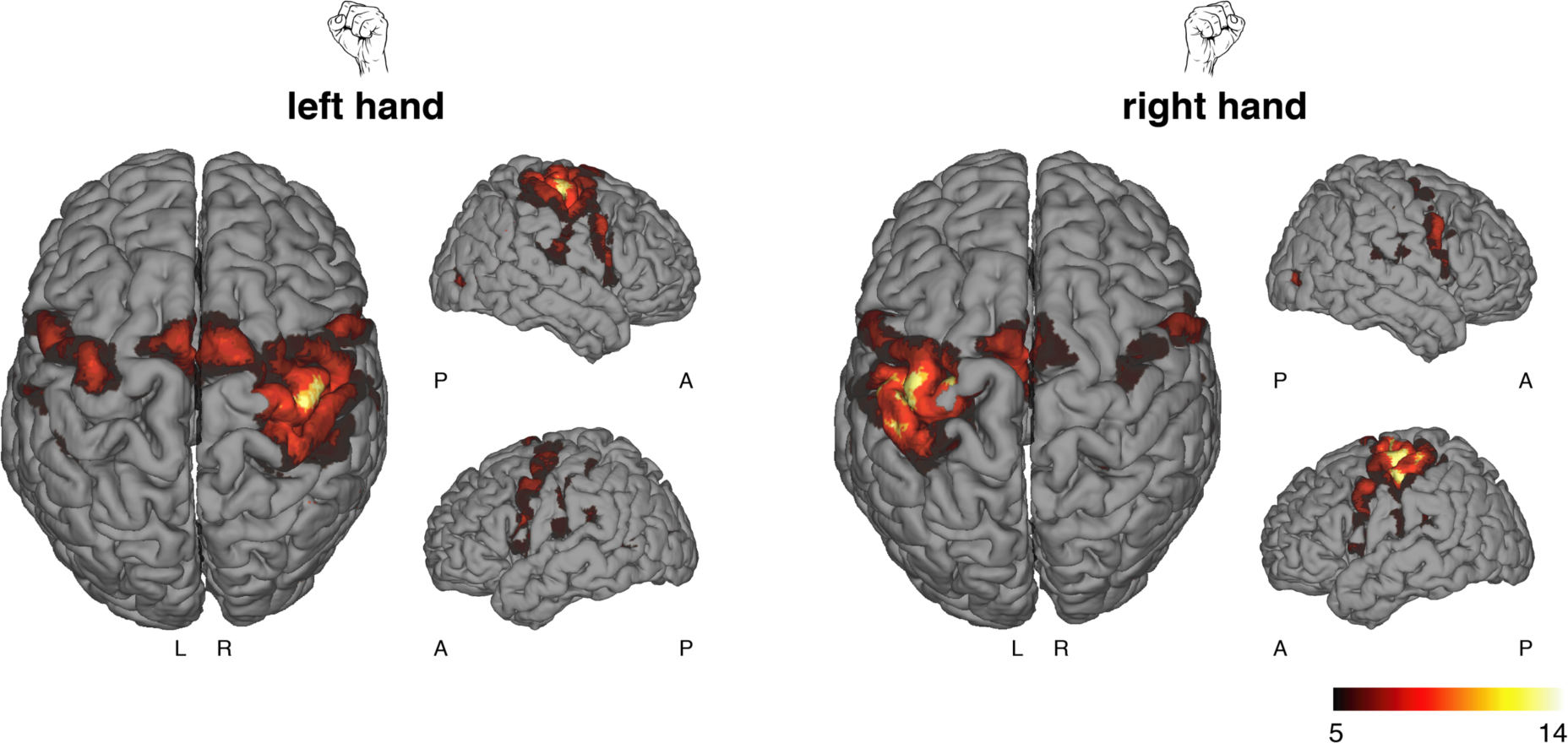
BOLD activation during visually synchronized unilateral hand movements at the group level (N=25). Left-hand (*left*) and right-hand fist closings (*right*) elicited activation in a distributed network, mainly lateralized to the contralateral hemisphere. Results are significant at a voxel-level threshold of *p*<0.05 (family-wise error (FWE)-corrected). Results were rendered onto the surface of an anatomical template volume. L = left hemisphere; R = right hemisphere; A = anterior; P = posterior.

### 3.2 Regression DCM constrained by anatomical connectivity

#### 3.2.1 Whole-brain effective connectivity during hand movements

Individual connectivity parameters were estimated using rDCM where, in a first step, the network architecture of the DCMs was informed by the structural connectome from the Brainnetome atlas (model 1; Figure 1A). Model inversion resulted in biologically plausible connectivity (Figure 3B, left) and driving input patterns (Figure 3B, right), suggesting pronounced functional integration in a widespread cortical network during visually paced unilateral hand movements. Consistent with our hypotheses (see Methods), rDCM revealed pronounced clusters of excitatory connections among motor and visual regions. Specifically, strong connections were observed among motor regions in the precentral (Brainnetome parcel name: *A4ul*) and postcentral gyrus (*A1/2/3ulhf*, *A2*), as well as the dorsal PMC (*A6cdl*) and the dorsal part of area 4 (*A4t*). Similarly, prominent functional integration was observed for the SMA (*A6m*) located in the superior frontal gyrus, as well as regions in the lateral occipital cortex, including the middle occipital gyrus (*mOccG*) and the motion-sensitive area (*V5/MT*). We also observed pronounced connections among regions in the parietal lobe (e.g., *A7c*, *A7m*, *A5m*), as well as excitatory connections from the parietal cortex to the visuomotor network highlighted above. Finally, connectivity was observed among frontal regions (e.g., *A8m*, *A6cvl*, *A44v*), as well as between frontal regions and all other components mentioned above. Overall, the majority of connections had positive weights (i.e., excitatory effects), which is consistent with the fact that our model describes changes of activity from baseline (i.e., activity induced by hand movements compared to rest). Furthermore, functional integration was strongest within hemispheres; however, pronounced interhemispheric connections were also observed, mainly among homotopic regions.

**Figure 3:**
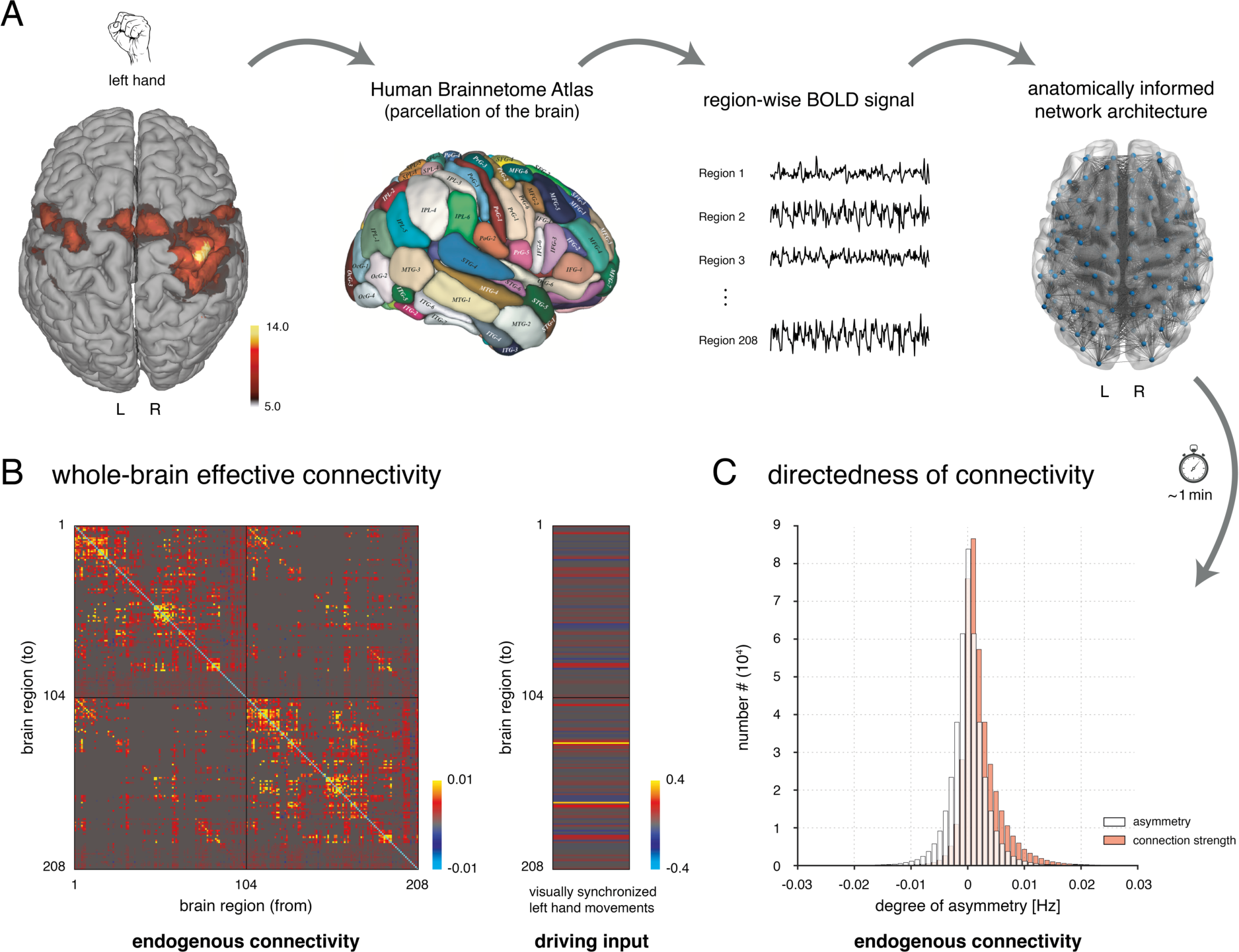
Whole-brain effective connectivity pattern underlying unilateral hand movements as assessed with rDCM when using structural connectivity to inform the network architecture. **(A)** For the given BOLD activation pattern during visually synchronized hand movements, the Human Brainnetome atlas^51^ was used as a whole-brain parcellation scheme. Region-wise BOLD signal time series were extracted for each participant individually as the principal eigenvariate and entered effective connectivity analyses using rDCM. **(B)** Posterior parameter estimates for connections (*left*) and driving inputs (*right*) during left-hand movements. **(C)** Histogram of asymmetry between the afferent (incoming) and efferent (outgoing) part of reciprocal connections (*white*). This suggests that the asymmetry was comparable in magnitude with the connection strengths themselves (*red*). Note that connectivity and driving input parameters represent rate constants and are thus given in Hz.

With regard to driving inputs (representing visually synchronized hand movements), we observed strong excitatory inputs to the motor and visual regions mentioned above (Figure 3B, right). Driving inputs to motor-related regions were stronger for nodes in the contralateral as compared to the ipsilateral hemisphere.

Notably, the directedness of connectivity estimates obtained by rDCM is demonstrated by the fact that, for the present dataset, there are pronounced asymmetries between the afferent (incoming) and efferent (outgoing) parts of reciprocal connections. Specifically, differences in the strengths of afferent and efferent connections were comparable in magnitude with the connection strengths themselves (Figure 3C), indicating a marked degree of directedness in the inferred connectivity patterns.

#### 3.2.2 Mirror symmetry of left- and right-hand movements

Next, we investigated the effect of the hand movement condition (i.e., left vs. right hand) by testing, for each parameter, whether there was a significant difference between left- and right-hand fist closings (two-sided paired *t*-test). We found the expected mirror-symmetric pattern, with connections in the left hemisphere being increased during right-hand movements and, vice versa, connections in the right hemisphere being increased during left-hand movements (Figure 4). These effects were highly specific in that only connections among sensorimotor areas showed significant hemispheric differences (*p*<0.05, false discovery rate (FDR)-corrected for multiple comparisons across the 16,868 free parameters). Specifically, as expected for the task we used, we found increased intrahemispheric connectivity among regions in the contralateral precentral (M1 (*A4ul*), dorsal PMC (*A6cdl*)) and postcentral gyrus (SM1 (*A1/2/3ulhf*, *A2*)). Furthermore, intrahemispheric connectivity was increased among the contralateral SMA (*A6m*) and M1 and SM1. Finally, rDCM revealed increased interhemispheric connections among SMA and M1 and SM1 (although this was not significant for the connections between right SMA and left pre- and postcentral gyrus when correcting for multiple comparisons).

**Figure 4:**
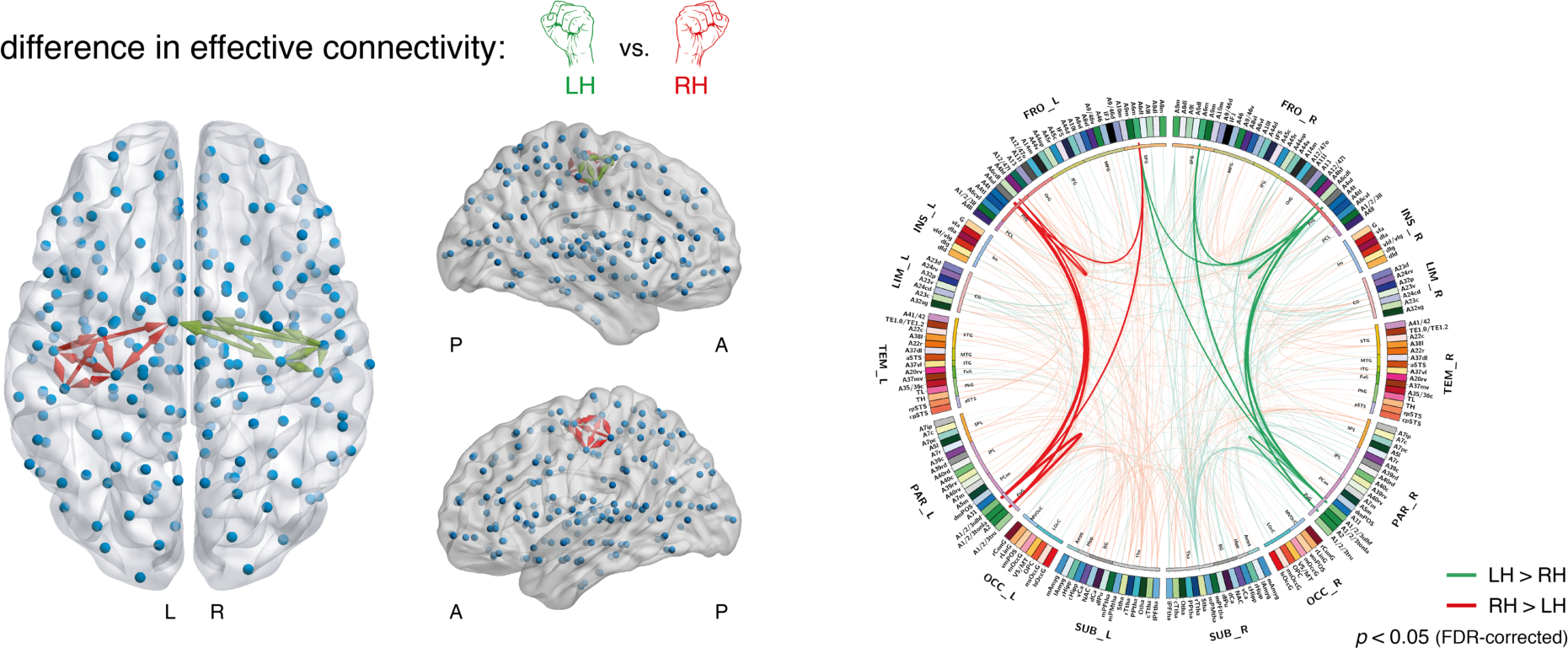
Mirror symmetry of the effect of hand movement condition (i.e., left vs. right hand) in the motor network as assessed with rDCM when using an anatomically informed (fixed) network architecture. The differential effect of hand movement condition was graphically projected onto a whole-brain volume (*left*). Green arrows indicate connections that were significantly increased during left-hand movements as compared to right-hand movements; red arrows indicate connections that were significantly increased during right-hand movements compared to left-hand movements (*p*<0.05, FDR-corrected for multiple comparisons). Note that edges in this graphical representation are directed. L = left hemisphere; R = right hemisphere; A = anterior; P = posterior. Results can also be inspected when graphically rendered as a connectogram (*right*). Solid lines represent the connections that showed a significant effect of the hand movement condition (*p*<0.05, FDR-corrected). Lines with faded colors represent the subsequent 500 connections with the strongest differential effect (highest absolute T values of the two-sided paired *t*-test). The labels on the outermost ring show the anatomical lobe for each of the nodes: frontal, insula, limbic, temporal, parietal, occipital, and subcortical. For each brain region defined by the Brainnetome atlas, an abbreviation and color are defined. Inside the parcellation ring, connections showing a significant effect of the hand movement condition are displayed as edges, with the color code defined as above (i.e., green = LH>RH, red = RH>LH).

#### 3.2.3 Benefit of informing network architecture with structural information

One might wonder whether utilizing the structural connectome from the Brainnetome atlas^51^ to inform the network architecture of the whole-brain DCMs was beneficial for explaining the observed fMRI data. To this end, we constructed two alternative networks: Model 2 (Figure 1B) represents a randomly permuted version of the Brainnetome structural connectome, and model 3 (Figure 1C) assumes a fully (all-to-all) connected network where all regions are linked via reciprocal connections. Since functional integration in the brain is constrained (but not fully determined) by anatomical connections^11, 53^, one would expect that effective connectivity analyses benefit from including tractography-based measures.

We used random effects Bayesian model selection (BMS)^54^ to compare the competing whole-brain models based on their log model evidence (approximated by negative free energy). We found decisive evidence that the anatomically informed model 1 was the winning model with a protected exceedance probability of 1. This illustrates clearly that models of whole-brain effective connectivity profit from structural connectivity measures derived from probabilistic tractography of DWI data. This is consistent with previous work in conventional (small-scale) DCMs that highlight the benefit of anatomically informed priors^55, 56^.

### 3.3 Regression DCM with sparsity constraints

#### 3.3.1 Whole-brain effective connectivity during hand movements

Next, we asked whether sensible whole-brain effective connectivity patterns could also be obtained in the absence of any *a priori* assumptions about the network’s architecture. For this, rDCM with embedded sparsity constraints was used to prune, for each participant individually, a fully connected model containing over 43,000 free connectivity parameters (Figure 5A).

**Figure 5:**
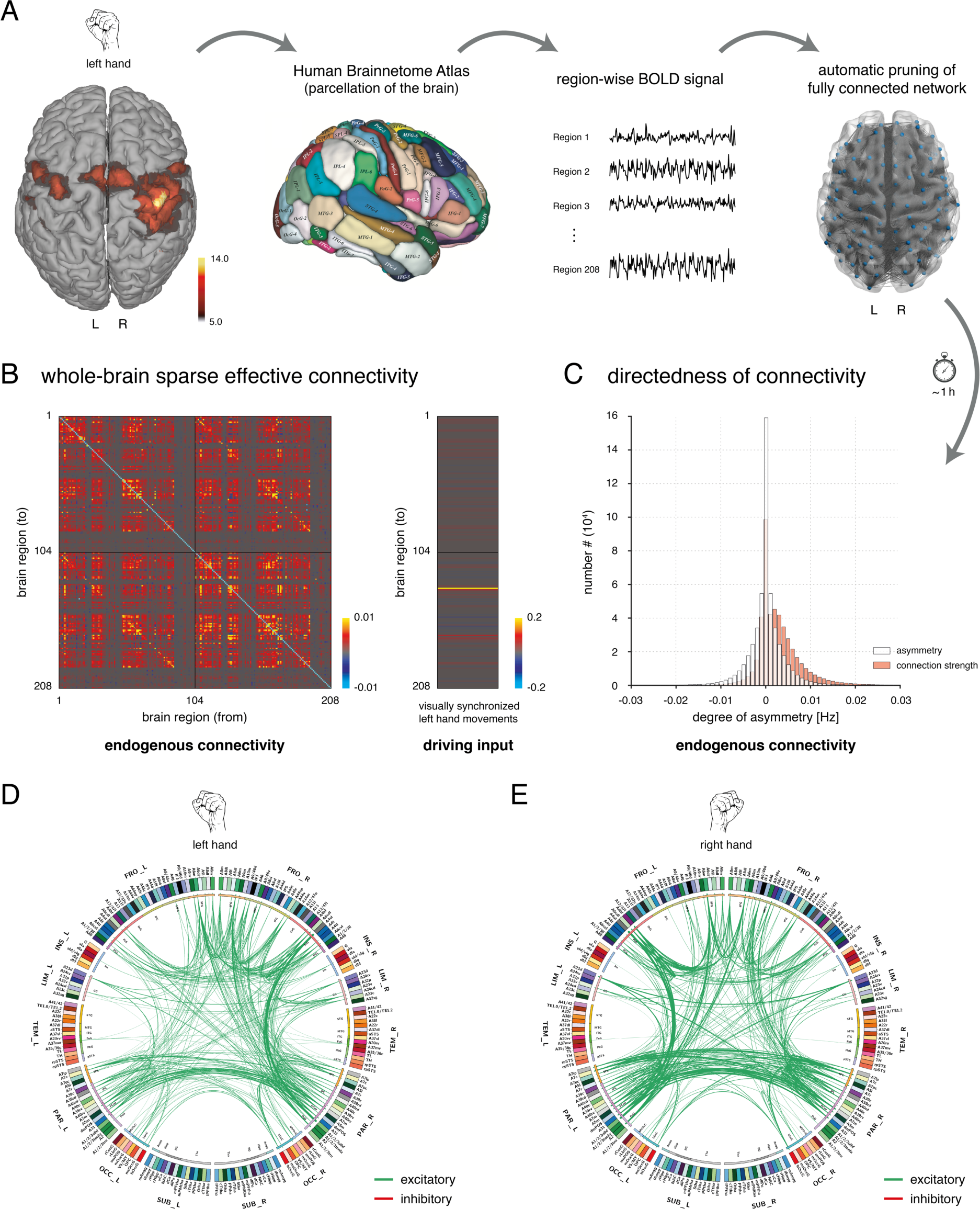
Sparse whole-brain effective connectivity pattern underlying unilateral hand movements as assessed with rDCM when embedded sparsity constraints were used to prune a fully (all-to-all) connected network. **(A)** For the given BOLD activation pattern during visually synchronized hand movements, the Human Brainnetome atlas^51^ was used as a whole-brain parcellation scheme. Region-wise BOLD signal time series were extracted for each participant individually as the principal eigenvariate and entered effective connectivity analyses using rDCM. A fully connected network was assumed and then pruned to an optimal (with respect to the negative free energy) degree of sparsity during model inversion. **(B)** Posterior parameter estimates for connections (*left*) and driving inputs (*right*) during left-hand movements. **(C)** Histogram of asymmetry between the afferent (incoming) and efferent (outgoing) part of reciprocal connections (*white*). This suggests that the asymmetry was comparable in magnitude with the connection strengths themselves (*red*). **(D)** Lines represent the 500 connections with the strongest effect for left-hand movements (i.e., highest absolute T value of the two-sided one-sample *t*-test for LH vs. baseline) **(E)** and right-hand movements (i.e., RH vs. baseline). The labels on the outermost ring show the anatomical lobe for each of the nodes: frontal, insula, limbic, temporal, parietal, occipital, and subcortical. For each brain region defined by the Brainnetome atlas, an abbreviation and color are defined. L = left hemisphere; R = right hemisphere.

Model inversion again revealed pronounced functional integration in a widespread network (Figure 5B). In brief, as expected and consistent with the anatomically constrained analysis, the sparse connectivity patterns revealed pronounced clusters of excitatory connections among regions in the motor (e.g., *A4ul*, *A6cdl*) and somatosensory cortex (e.g., *A1/2/3ulhf*, *A2*), occipital lobe (e.g., *mOccG*, *V5/MT*), as well as parietal cortex (e.g., *A39rd/rv*, *A40rd*/*rv*, *A7m*), and frontal lobe (e.g., *A6vl*, *A8vl*, *A44v*). Again, the majority of connections were of positive sign (i.e., excitatory), reflecting the fact that our model describes activity changes relative to rest. With regard to driving inputs, excitatory effects were observed for regions in the contralateral precentral (*A4ul*, *A4t*, *A6cdl*) and postcentral gyrus (*A1/2/3ulhf*, *A2*). Additionally, we found driving inputs to SMA (*A6m*) and visual regions, including the middle occipital gyrus (*mOccG*) and the motion-sensitive area (*V5/MT*).

As for the tractography-guided application of rDCM, we tested whether the sparse effective connectivity estimates showed asymmetries between afferent and efferent connections. As above, differences in the strength between afferent and efferent connections were comparable in magnitude with the connection strengths themselves (Figure 5C). This demonstrates that rDCM estimates displayed a marked degree of directedness also when embedded sparsity constraints were used.

For rDCM under sparsity constraints, which in contrast to the anatomically informed analysis does not rely on a symmetric structural connectome, it is instructive to inspect the top 500 connections for both left- and right-hand movements (Figure 5D-E). This plot illustrates the expected contralateral lateralization of the connectivity pattern – in particular, for connections among pre- and postcentral gyrus, as well as for connections from superior frontal gyrus (e.g., *A6m*) and parietal regions to premotor and motor regions. Finally, for both left- and right-hand movements, one can observe strong interhemispheric connections that were most pronounced among homotopic areas in frontal and parietal cortex.

#### 3.3.2 Mirror symmetry of left- and right-hand movements

As for the anatomically informed rDCM analysis, we explicitly assessed the effect of hand movement condition (i.e., left vs. right hand). Again, we found the expected mirror-symmetric pattern, with connections in the left hemisphere being increased during right-hand movements and, vice versa, connections in the right hemisphere being increased during left-hand movements (Figure 6). Significant effects (*p*<0.05, FDR-corrected for multiple comparisons across the 43,472 free parameters) were again constrained to connections among sensorimotor regions. We observed an effect of the hand movement condition for the intrahemispheric connections among M1 (*A4ul*), SM1 (*A1/2/3ulhf*, *A2*), and SMA (*A6m*).

**Figure 6:**
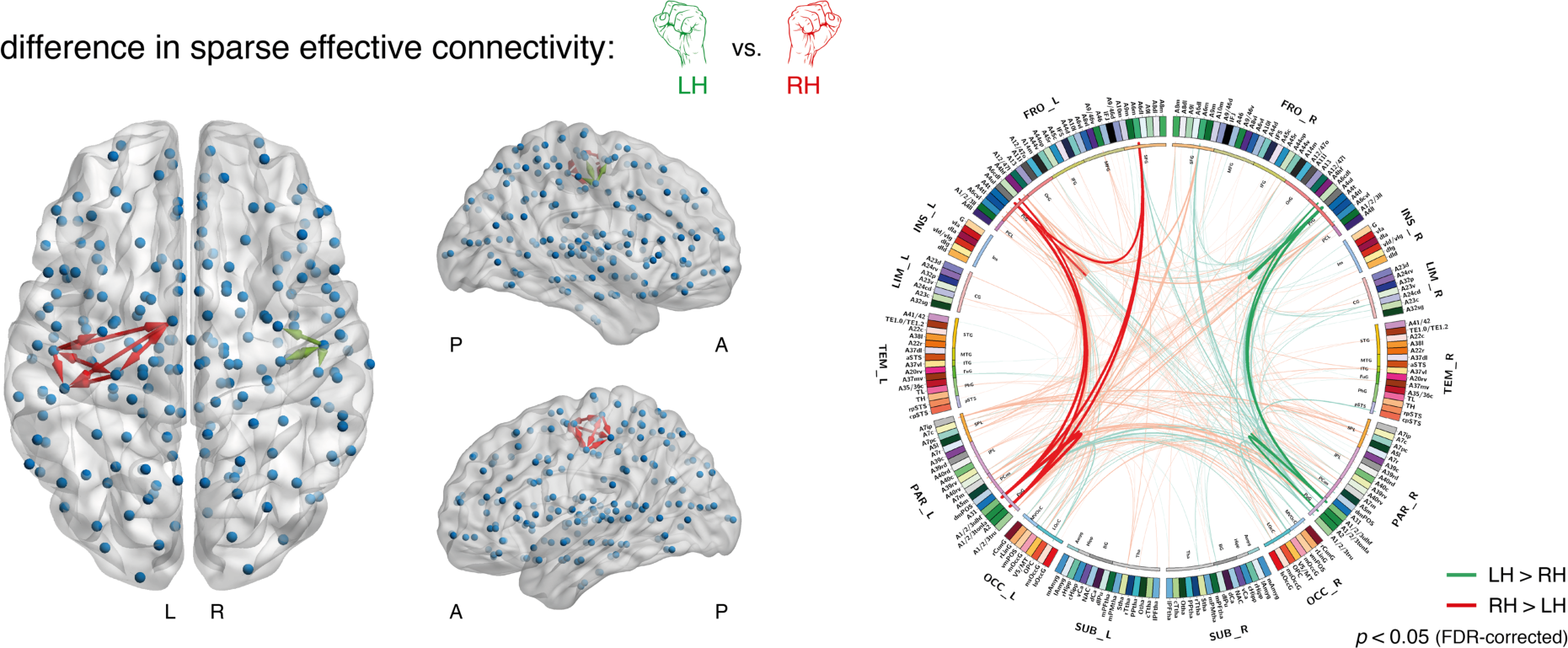
Mirror symmetry of the effect of hand movement condition (i.e., left vs. right hand) in the motor network as assessed using rDCM with embedded sparsity constraints to prune a fully (all-to-all) connected network. The differential effect of hand movement condition was graphically projected onto a whole-brain volume (*left*). Green arrows indicate connections that were significantly increased during left-hand movements as compared to right-hand movements; red arrows indicate connections that were significantly increased during right-hand movements compared to left-hand movements (*p*<0.05, FDR-corrected for multiple comparisons). Note that edges in this graphical representation are directed. L = left hemisphere; R = right hemisphere; A = anterior; P = posterior. Results can also be inspected when graphically rendered as a connectogram (*right*). Solid lines represent the connections that showed a significant effect of the hand movement condition (*p*<0.05, FDR-corrected). Lines with faded colors represent the subsequent 500 connections with the strongest differential effect (highest absolute T values of the two-sided paired *t*-test). The labels on the outermost ring show the anatomical lobe for each of the nodes: frontal, insula, limbic, temporal, parietal, occipital, and subcortical. For each brain region defined by the Brainnetome atlas, an abbreviation and color are defined. Inside the parcellation ring, connections showing a significant effect of the hand movement condition are displayed as edges, with the color code defined as above (i.e., green = LH>RH, red = RH>LH).

#### 3.3.3 Graph-theoretical analyses

In a next step, we applied graph-theoretical measures^53^ to the sparse whole-brain effective connectivity patterns underlying unilateral hand movements. Specifically, using graph theory, we intended to corroborate the pivotal role of motor regions in the pre- and postcentral gyrus during our task, as well as the known hemispheric lateralization of the network. To this end, we chose graph-theoretical measures that capture the importance/relevance of each node and that have frequently been used in the field of connectomics: “betweenness centrality” and “node strength (in & out)”. In brief, betweenness centrality is the fraction of all shortest paths in the network that contain a given node, whereas node strength refers to the sum of weights of all links connected to a node. We tested whether graph-theoretical measures would more faithfully reflect known functional properties of the motor system when applied to directed as compared to undirected connectivity measures. Graph-theoretical measures were computed using the Brain Connectivity toolbox^57^.

Figure 7 shows the betweenness centrality for each of the 208 parcels from the Brainnetome atlas (projected onto a whole-brain volume) for left- and right-hand movements. The expected contralateral dominance of the motor regions is clearly visible: For left-hand movements, the node with the highest betweenness centrality was right M1; whereas, for right-hand movements, left M1 showed one of the highest betweenness centrality scores (Figure 7A-B). We also found high betweenness centrality scores during unilateral hand movements in regions located in the contralateral somatosensory cortex (*A1/2/3ulhf*, *A2*). Furthermore, high betweenness centrality in both left and right hemisphere, regardless of the hand movement condition, was observed in the medial area 7 (*A7m*), which represents the visuospatial/-motor part of the precuneus.

**Figure 7:**
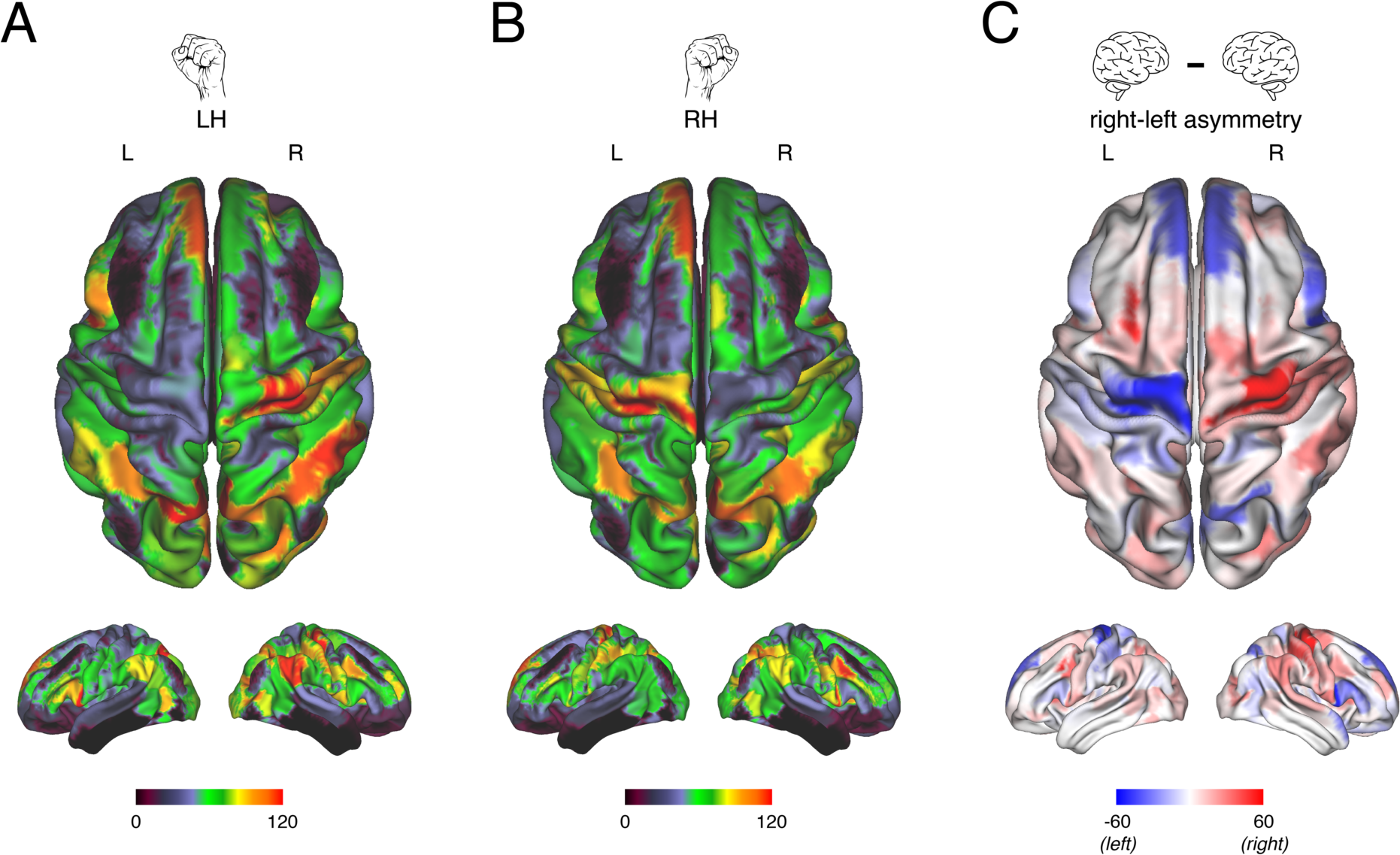
Graph-theoretical analysis of the whole-brain effective connectivity patterns underlying unilateral hand movements as inferred using rDCM when embedded sparsity constraints were used to prune a fully (all-to-all) connected network. Betweenness centrality was evaluated for each parcel of the Human Brainnetome atlas for **(A)** left-hand and **(B)** right-hand fist closings and then graphically projected onto a whole-brain volume. **(C)** Hemispheric asymmetries in betweenness centrality were assessed by evaluating the difference in betweenness centrality for homotopic parcels in the left and right hemisphere. Positive values indicated higher betweenness centrality in the right hemisphere (*red*), whereas negative values indicated higher betweenness centrality in the left hemisphere (*blue*). Results for left-hand movements are presented in the right hemisphere, whereas results for right-hand movements are presented in the left hemisphere. Again, this clearly illustrates the mirror symmetry of the motor network in the pre- and postcentral gyrus. Betweenness centrality for directed and weighted adjacency matrices was computed using the Brain Connectivity toolbox^57^, which is freely available (https://sites.google.com/site/bctnet/). Betweenness centrality values for each parcel were visualized using the Human Connectome Workbench, also publicly available (https://www.humanconnectome.org/software/connectome-workbench). L = left hemisphere; R = right hemisphere; LH = left hand; RH = right hand.

Hemispheric differences in betweenness centrality revealed the expected mirror-symmetric pattern within motor-related regions in the precentral (*A4ul*) and postcentral gyrus (*A1/2/3ulhf*, *A2*). Specifically, hemispheric asymmetry in these regions depended strongly on the hand movement condition (Figure 7C): betweenness centrality was higher in the right hemisphere during left-hand movements, and higher in the left hemisphere during right-hand movements.

Notably, the mirror symmetry of functional integration during left- and right-hand movements was not a global finding, but was specific to the motor network. In contrast, regions in the frontal (e.g., *A6dl*, *A46*, *A8vl*, *A44d*) and parietal lobe (e.g., *A7r*) showed higher betweenness centrality in the right hemisphere, regardless of the hand movement condition. Furthermore, regions in the occipital lobe, such as the primary visual cortex in the occipital polar cortex (*OPC*) and the motion sensitive area *V5/MT*, did not show marked hemispheric asymmetries, consistent with the central visual stimulation during both hand movement conditions.

For node strength, results were highly consistent with the pattern observed for betweenness centrality, again highlighting the contralateral dominance of motor regions and the expected mirror symmetry of the network for left- and right-hand movements (Supplementary Figure S2).

#### 3.3.4 Sparsity constraints vs anatomical constraints

In a final step, we compared the two general modes of operation for rDCM: fixed network architecture informed by a structural connectome (anatomical constraints) versus pruning a fully connected whole-brain model (sparsity constraints). First, one can observe that the effective connectivity pattern under anatomical constraints (Figure 3B) is not dissimilar to the product of the fixed Brainnetome structural connectome serving as prior (Figure 1A) and the inferred pattern under sparsity constraints (Figure 5B), which intuitively is plausible. Second, since rDCM provides a principled measure of model goodness, the log model evidence, one can use BMS to ask which mode provided a better explanation of the data. Random effects BMS indicated that the model with anatomically informed (fixed) network architecture was superior with a protected exceedance probability of 1. This suggests that – in this case – exploiting available anatomical information to inform the architecture of the model was clearly beneficial.

### 3.4 Computational burden

Concerning computational efficiency, running model inversion on a single processor core (without parallelization) on the Euler cluster at ETH Zurich (https://scicomp.ethz.ch/wiki/Euler), rDCM took on the order of a minute or less when assuming structurally fixed connectivity and input structure. More specifically: for models 1 and 2 (16,868 free parameters), model inversion took around 20s, whereas for model 3 (43,472 free parameters), model inversion took roughly 100s.

Using sparsity constraints to prune fully connected networks is computationally more demanding: on average (across participants), rDCM took roughly 4h on a single processor core to infer sparse connectivity patterns under a given *p*^i^_0_ value. This compares favorably to other methods of large-scale effective connectivity, like cross-spectral DCM, for which 21-42h of run-time on a high-performance computing cluster for a network with 36 regions and 1,260 connections has been reported^30^.

Notably, these run-times were obtained using a language not optimized for speed (Matlab) nor without any effort to speed the code up by parallelization. The latter is a straightforward and powerful option to further enhance the efficiency of rDCM^33^. This is due to the mean field approximation in rDCM which allows applying the VB update equations to each region independently. Specifically, when using 16 processor cores in parallel, the above run-time for inferring sparse effective connectivity patterns could be reduced to around 40min on average. The values reported here should only be treated as a rough indication, as run-times will depend on the specific hardware used.

### 3.5 Comparison to undirected measures of brain connectivity

In a final step, we compared the whole-brain effective connectivity estimates with measures of functional connectivity, which represent the current standard in human connectomics. For this, we computed for each participant Pearson correlation coefficients between exactly the same 208 BOLD signal time series as used in the rDCM analysis. Pearson correlations arguably represent the simplest and most widely used measures of functional connectivity. In contrast to the Bayesian framework of rDCM, Pearson correlations are not subject to any regularization, which has advantages and disadvantages: they might be more sensitive for detecting functional coupling, but are also very sensitive to measurement noise^58^.

Functional connectivity patterns for the unilateral hand movements were qualitatively similar to the effective connectivity patterns obtained using rDCM: we observed coupling among motor (i.e., precentral, SMA), visual (occipital), somatosensory/proprioceptive (postcentral, parietal) and frontal regions (Figure 8A). However, in contrast to effective connectivity (cf. Figure 3C and 5B), functional connectivity does not afford any information on the directionality of influences, leading to symmetric connectivity matrices.

**Figure 8:**
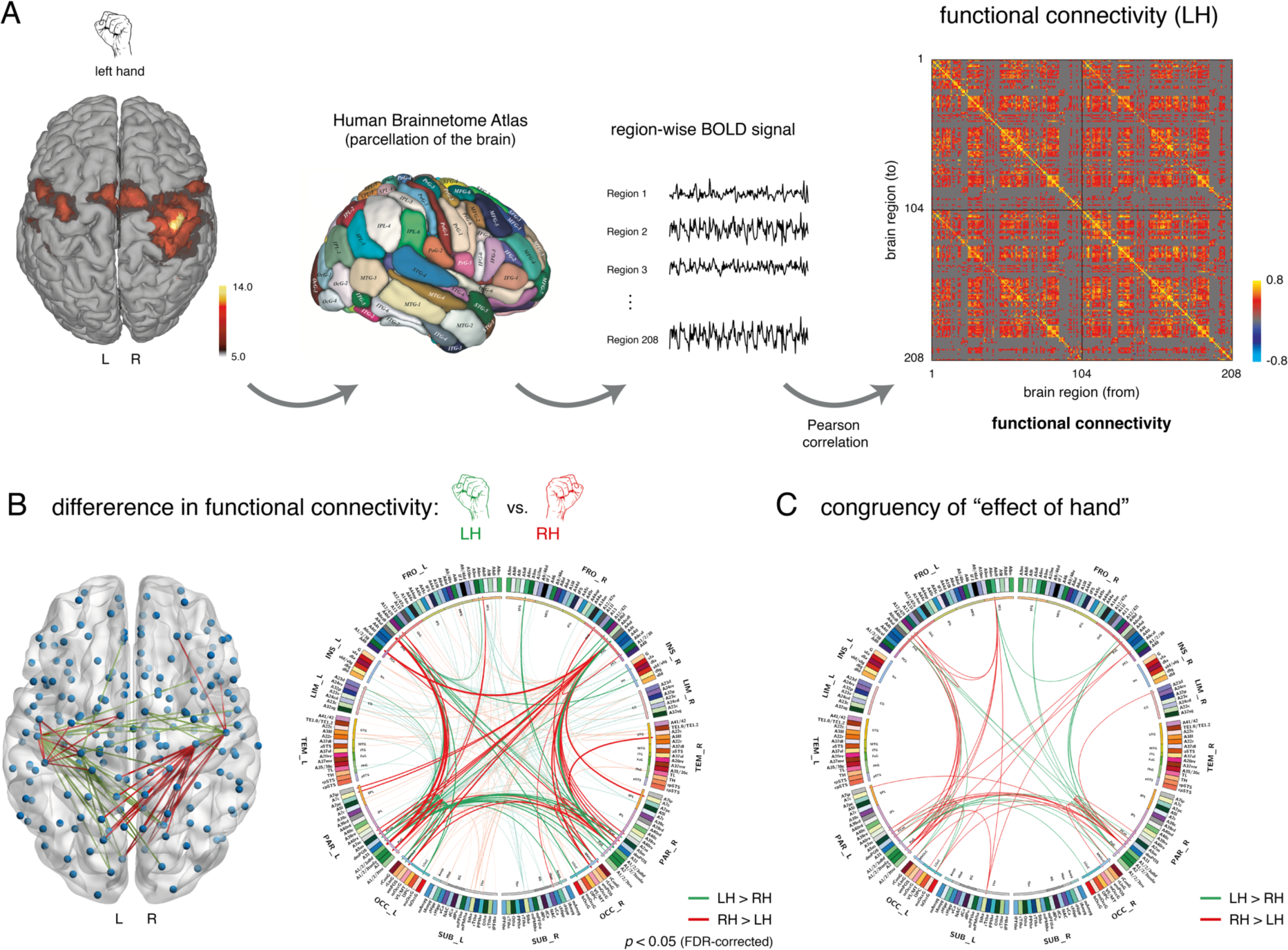
Functional connectivity pattern underlying unilateral hand movements as assessed using the Pearson correlation coefficient among the same BOLD signal time series as utilized for the rDCM analysis. **(A)** For the given BOLD activation pattern during visually synchronized hand movements, the Human Brainnetome atlas^51^ was used as a whole-brain parcellation scheme. Region-wise BOLD signal time series were extracted for each participant individually as the principal eigenvariate and entered functional connectivity analyses using the Pearson correlation coefficients. To allow for comparability with rDCM results, the functional connectivity matrix is thresholded such that the sparsity of the matrix resembles the sparsity of the structural connectome from the Brainnetome atlas. **(B)** Differential effect of hand movement condition on functional connectivity, projected onto a whole-brain volume (*left*). Green lines indicate connections that were significantly increased during left-hand movements as compared to right-hand movements; red lines indicate connections that were significantly increased during right-hand movements compared to left-hand movements (*p*<0.05, FDR-corrected). L = left hemisphere; R = right hemisphere; LH = left hand; RH = right hand. Results can also be rendered as a connectogram (*right*). Solid lines represent significant differential effects (*p*<0.05, FDR-corrected), faded colors represent the 500 connections with the next highest absolute T values of the two-sided paired *t*-test. The labels on the outermost ring show the anatomical lobe for each of node: frontal, insula, limbic, temporal, parietal, occipital, and subcortical. Next, abbreviation and color for each region are shown. **(C)** Connections showing a congruent effect of hand movement condition on the functional and effective connectivity estimates. Congruency was established by logical AND of the connectomes in Figures 4 and 8B, where rDCM estimates were first converted into undirected connections before binarizing all connections as having positive and negative strengths. Lines indicate those of the top 500 connections of the functional and effective connectivity patterns that showed the same differential hand movement effect (i.e., LH>RH or RH>LH).

We then tested for the differential effect of the hand movement condition (i.e., left vs. right hand) using two-sided paired *t*-tests (*p*<0.05, FDR-corrected) after Fisher r-to-z transformation of the correlation coefficients. Consistent with rDCM, intrahemispheric functional connectivity among M1 (*A4ul*) and SM1 (*A1/2/3ulhf*, *A2*) of the contralateral hemisphere was increased (Figure 8B). However, functional connectivity did not show the expected mirror-symmetric pattern within the motor network as clearly as in the case of rDCM: Various connections within the motor network (and beyond) showed the opposite effect, resulting in a more ambiguous pattern. Furthermore, no significant effect could be observed for connections between SMA (*A6m*) and regions in the pre- and postcentral gyrus when correcting for multiple comparisons. This was slightly unexpected given the prominent role of the SMA in the initiation of voluntary hand movements^41, 47^.

To compare functional and effective connectivity estimates more directly, we computed a congruence map between functional connectivity and rDCM, covering the 500 connections with the strongest effect of the hand movement condition (for details, see legend to Figure 8C). While the majority of connections did not overlap between the two methods, those connections that showed strong differences between hand conditions (mainly connections among motor-related regions) displayed the same sign for functional connectivity and rDCM estimates (Figure 8C). This indicates that, at the level of undirected connections, functional and effective connectivity estimates are qualitatively compatible for those connections that are expected to be most relevant for the task.

We repeated the graph-theoretical analyses by evaluating betweenness centrality and node strength for the undirected functional connectivity patterns. In contrast to effective connectivity, functional connectivity did not show the expected pattern of betweenness centrality. Specifically, motor-related regions in the contralateral precentral (*A4ul*) and postcentral gyrus (*A1/2/3ulhf*, *A2*) did not yield high betweenness centrality scores (Figure 9A-B), in contradiction to their established role during unilateral hand movements. Furthermore, when testing for hemispheric differences in betweenness centrality, we did not observe the expected mirror symmetry in the motor network (Figure 9C). Similarly, node strength did not capture the importance of motor-related regions and yielded counterintuitive hemispheric asymmetries (Supplementary Figure S3), with a node strength pattern of motor-related regions opposite to what one would expect. This result may have been driven by connections between motor and more occipital regions that showed unexpected effects of hand in the functional connectivity analyses (Figure 8B). These unexpected findings may reflect the known sensitivity of correlation-based functional connectivity estimates to measurement noise^58^.

**Figure 9:**
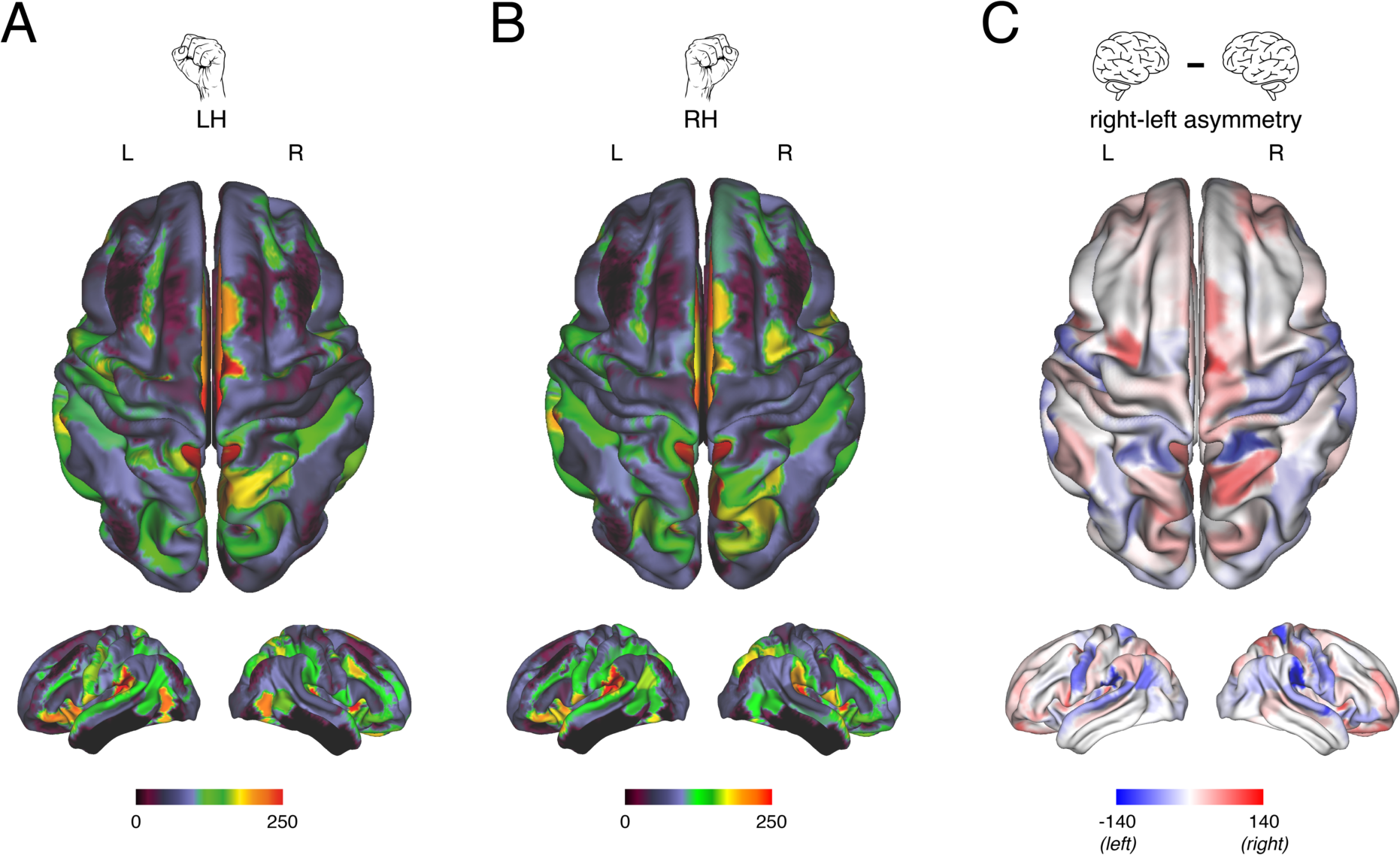
Graph-theoretical analysis of the whole-brain functional connectivity patterns during unilateral hand movements as assessed using the Pearson correlation coefficient. Betweenness centrality was evaluated for each parcel of the Human Brainnetome atlas for **(A)** left-hand and **(B)** right-hand fist closings and then graphically projected onto a whole-brain volume. **(C)** Hemispheric asymmetries in betweenness centrality, with positive values indicating higher betweenness centrality in the right hemisphere (*red*) and negative values indicating higher betweenness centrality in the left hemisphere (*blue*). Results for left-hand movements are presented in the right hemisphere, whereas results for right-hand movements are presented in the left hemisphere.

## 4 DISCUSSION

In this paper, we assessed the construct validity of regression DCM (rDCM) for inferring whole-brain effective connectivity patterns from fMRI data. Using a hand movement dataset, we demonstrated that rDCM can infer plausible effective connectivity patterns in a network comprising over 200 regions and 40,000 free parameters. Furthermore, we applied graph-theoretical measures to the whole-brain effective connectivity patterns and demonstrate that they capture the expected pivotal role of motor-related regions, as well as the hemispheric asymmetries of the network.

In brief, rDCM identified pronounced functional integration among key components of the motor network – e.g., M1, SM1, and SMA. Furthermore, when testing for effects of the hand movement condition (i.e., left vs. right hand), we found the expected mirror-symmetric pattern: connections among key motor regions in the left hemisphere were increased during right-hand movements and, vice versa, connections in the right hemisphere were increased during left-hand movements. This pattern could not only be obtained when structural connectivity data were used to inform the network architecture of the whole-brain DCMs, but even in the case of complete absence of *a priori* assumptions about the network’s architecture by automatically pruning fully connected graphs to an optimal degree of sparsity.

However, our method also failed to detect a characteristic of the motor system that has been reported previously: interhemispheric inhibition of the ipsilateral M1 by the contralateral M1 during unilateral hand movements^59^. This may be due to the fact that hand movements of different conditions were separated into two scanning sessions, potentially rendering interhemispheric inhibition less critical as in paradigms that alternate between the two conditions^41^.

We further demonstrated the application of graph-theoretical measures to the inferred whole-brain effective connectivity patterns. Specifically, we show that measures that capture the relevance of a network node, i.e., betweenness centrality and node strength, correctly identify motor-related regions in the pre- and postcentral gyrus as key components of the network and show the expected hemispheric asymmetry^60, 61^. Furthermore, our graph-theoretical analyses are consistent with known functional characteristics of the human brain, including the relevance of precuneus in directing spatial attention during preparation and execution of motor actions^62–64^ and its role as a central “small-world network” hub^53^. Similarly, our analyses revealed the expected right-hemispheric lateralization of the fronto-parietal network underlying visuospatial attention^65, 66^.

Recently, graph theory has found widespread application in neuroscience and has provided valuable insights into the organization of the brain^53, 57, 67^. However, to render graph-theoretical approaches, and connectomics in general, meaningful for understanding organizational principles in large-scale networks, individual connection estimates need to be neurobiologically interpretable. This is not the case for currently used standard measures of connectivity in humans, such as DWI-derived structural connectivity and fMRI-based functional connectivity. These measures are undirected and do not capture functional asymmetries of reciprocal connections^20^. Extending graph-theoretical approaches to effective (directed) connectivity may therefore be critical for exploiting the information provided by graph-theoretical indices and for providing a more faithful assessment of the network topology underlying brain dynamics. Indeed, our findings suggest that directed estimates of connectivity boost the explanatory power of network analyses: graph theory applied to rDCM estimates better match known functional roles of key motor regions than when undirected functional connectivity estimates are used.

We would have liked to report a comparison between rDCM and measures of directed functional connectivity, like multivariate Granger causality (GC)^68^. However, for the data used here, GC estimates did not show convergence, probably due to issues like TR and the relatively short scanning time (i.e., low number of data points per region)^68, 69^. By contrast, the feasibility of obtaining meaningful estimates by rDCM underscores its potential suitability for clinical applications, where long scanning sessions are usually not possible.

Our results suggest that rDCM confers important practical advantages for human connectomics and network neuroscience. Several strengths and innovations are worth highlighting. First, rDCM provides different modes of operation for deriving directed connectivity fingerprints: it can exploit subject-specific anatomical connectivity information for constraining inference; alternatively, when no such information is available, rDCM can infer optimally sparse representations of whole-brain networks. Second, by introducing sparsity constraints, rDCM circumvents the need for arbitrary thresholding of connectivity matrices. Instead, rDCM yields an optimal degree of sparsity by maximizing the model evidence. Finally, rDCM is computationally highly efficient with run-times on the order of minutes per subject. This efficiency renders rDCM a promising tool for clinical applications but also for time-consuming analyses of large-scale datasets like the Human Connectome Project^70^. These developments provide exciting new opportunities for moving human connectomics and network neuroscience towards directed measures of functional integration. Furthermore, rDCM may find useful application in the emerging fields of Computational Psychiatry and Computational Neurology where computational readouts of directed connectivity in whole-brain networks are of major relevance^35, 71, 72^.

Despite these strengths, our study is also subject to limitations. First, the Brainnetome atlas does not cover the cerebellum^51^, which plays an important role in preparation and execution of motor actions^40^. This is similar to most other state-of-the-art whole-brain parcellation schemes, like the Human Connectome Project parcellation (HCP MMP 1.0), which are equally restricted to cortical regions. Other parcellation schemes, such as the Automated Anatomical Labeling atlas, do include the cerebellum but have other shortcomings. For the present analysis, we deliberately focused on the cortex and chose the Brainnetome atlas for three reasons: (i) the atlas is sufficiently fine-grained to allow for meaningful large-scale effective connectivity analyses among cortical regions, (ii) has been demonstrated to provide robust parcels across the population as assessed using cross-validation, and (iii) provides not only a parcellation of the brain but also tractography-based information on how these parcels are anatomically connected (which informed the network architecture in our initial rDCM analysis).

Second, rDCM is still in an early development stage and the current implementation is subject to methodological limitations^33, 34^. In particular, the biophysically plausible hemodynamic model in classical DCM was replaced with a fixed HRF. Consequently, rDCM presently does not capture variability in the BOLD signal across regions and individuals. In this work, we accounted for variability in latency and duration of the hemodynamic responses by including temporal and dispersion derivatives of the canonical HRF as confound regressors in the GLM^50^. Nevertheless, replacing the fixed HRF with a flexible hemodynamic model represents a major future development of rDCM.

It is worth highlighting that rDCM is not the only approach that aims to infer effective connectivity in large-scale networks. Alternative approaches include BNMs^28^ and cross-spectral DCMs^30^. BNMs combine biophysical mean-field models of the local neuronal dynamics with long-range connections informed by structural connectivity estimates. However, the complexity of these models renders parameter estimation computationally extremely challenging, restricting applications to relatively few free parameters^73–75^; but see ^32^ for notable progress in this area. A platform for constructing and applying BNMs to a variety of neuroimaging modalities is the Virtual Brain^76^.

Recently, a large-scale network model has been introduced that also enables inference on individual connectivity parameters^31^. Here, local dynamics of brain regions are described by an Ornstein-Uhlenbeck process. This model further differs from rDCM in that effective connectivity is not estimated within a Bayesian framework but by maximum likelihood, which does not enable automatic pruning of fully connected networks. For “resting-state” data, a variant of cross-spectral DCM has been proposed, where the effective number of free parameters is reduced by constraining the prior covariance matrix^30^. In contrast to rDCM, cross-spectral DCM explicitly captures regional variability in hemodynamic responses^77^. However, this increase in physiological realisms comes at the expense of non-optimal computational efficiency – resulting in run-times between 21-42h for a single DCM with 36 regions^30^. Hence, in its current implementation, cross-spectral DCM is unlikely to scale to whole-brain networks with hundreds of regions. In addition to cross-spectral DCM, alternative variants of large-scale connectivity analyses for “resting-state” fMRI data have recently been proposed that are inspired by rDCM and pursue a sparsity-inducing approach^78^.

In summary, in comparison to other methods for inferring effective connectivity in large-scale networks, rDCM provides estimates of the full posterior distributions of individual connection strengths in networks with hundreds of nodes, with run-times on the order of minutes on standard hardware. It allows for parallelization and scales gracefully with network size, an important property as methodological advances allow for increasingly fine-grained parcellations of human cortex^79^. Furthermore, its Bayesian formulation allows for a natural connection to subject-specific anatomical connectivity data (e.g., tractography). Finally, its speed and ability to prune whole-brain networks in the absence of anatomical connectivity information are important assets for clinical applications, potentially supporting time-sensitive clinical assessments with interpretable sparse whole-brain connectograms solely based on fMRI data.

## 5 CODE AND DATA AVAILABILITY

A Matlab implementation of the regression dynamic causal modeling (rDCM) approach is available as open source code in the **T**ranslational **A**lgorithms for **P**sychiatry-**A**dvancing **S**cience (TAPAS) Toolbox (www.translationalneuromodeling.org/software). Furthermore, following acceptance of this paper, we will publish the code for the analysis as well as the data used in this paper online as part of an online repository that conforms to the FAIR (Findable, Accessible, Interoperable, and Re-usable) data principles.

## ACKNOWLEDGEMENTS

This work was supported by the UZH Forschungskredit Postdoc (SF), the ETH Zurich Postdoctoral Fellowship Program and the Marie Curie Actions for People COFUND Program (SF), the René and Susanne Braginsky Foundation (KES), the Swiss National Science Foundation 320030_179377 (KES), and the University of Zurich (KES).

## 6 AUTHOR CONTRIBUTIONS

ZMM, KPP and KES conceptualized and designed the study, ZMM and LK performed the experiment, SF co-developed the modeling approach utilized in this study, SF and CTD preprocessed the data, SF analyzed the data, CTD performed a code review of the analysis pipeline, SF and KES discussed and interpreted the results, SF and KES wrote the manuscript, ZMM, CTD, LK, and KPP edited and approved the manuscript.

