## Supplementary Material for "Whole-brain estimates of directed connectivity for human connectomics"

#### Supplementary Material S1: Dynamic causal modeling

Dynamic causal modeling (DCM) is a generative modeling framework for inferring effective (directed) connectivity among latent (hidden) neuronal states from measured neuroimaging data[1]. Originally, DCM was introduced for functional magnetic resonance imaging (fMRI) data where changes in neuronal states can be described using a bilinear differential equation

$$\frac{dx}{dt} = \left( A + \sum_j B^{(j)} u_j \right) x + Cu \qquad \text{(S.1)}$$

In Eq. (S.1), changes in the neuronal states $x$ in time unfold as a function of the coupling among network nodes (endogenous connectivity $A$) and experimental manipulations $u$ (e.g., sensory stimulation, task demands) that perturb the neuronal system. Experimental manipulations can either modulate the endogenous connectivity to induce condition-specific changes in coupling strength (modulatory connectivity $B$) or directly affect the neuronal states in a brain region (driving inputs $C$). Effective connectivity parameters in DCM represent rate constants and are given in Hz. This neuronal state equation is coupled with a second set of differential equations that represents the hemodynamic model. The hemodynamic model rests on the Balloon-Windkessel model[2] which was augmented to account for neurovascular coupling[3]. This provides a biophysically informed model of how neuronal dynamics translate into region-wise blood oxygen level dependent (BOLD) signals. By assuming suitable (Gaussian) noise on the predicted BOLD signals, one obtains a probabilistic forward mapping (i.e., a likelihood function) that links latent neuronal states with measured fMRI data. The likelihood function is combined with prior distributions on model parameters and hyperparameters to give rise to a full generative model. Inversion of this generative model then rests on Variational Bayes under the Laplace assumption (VBL)[4,5]. Notably, since its initial introduction, various variants of DCM for fMRI have been proposed, including two-state DCM[6], nonlinear DCM[7], stochastic DCM[8,9], and spectral DCM[10,11]. For comprehensive reviews of DCM for fMRI, see refs. [9,12].

### Figures

#### Supplementary Figure S1

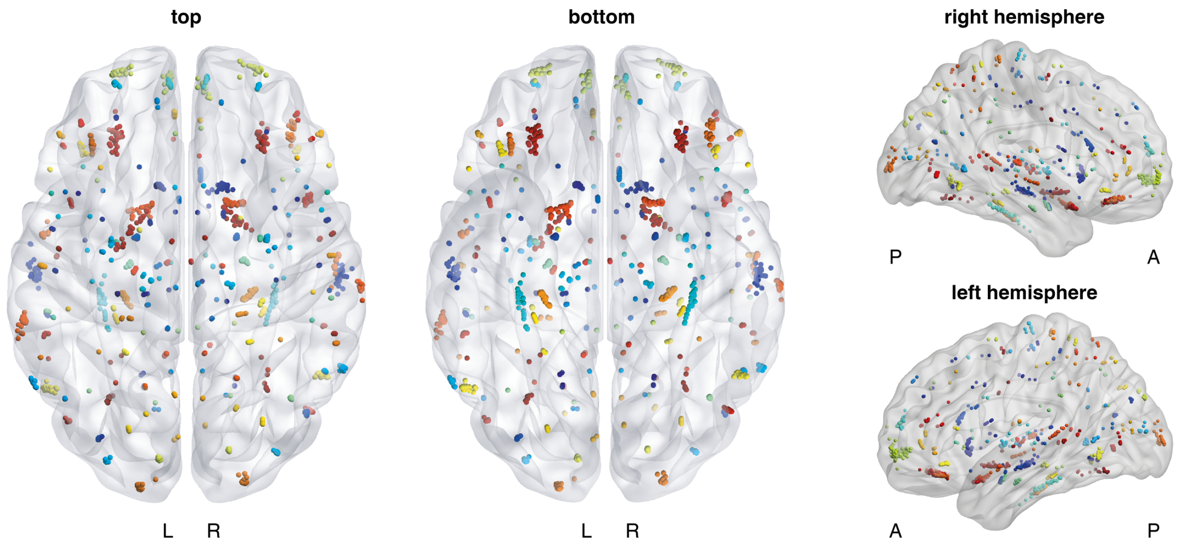


**Supplementary Figure S1:** Individual locations of each parcel from the Human Brainnetome atlas[13], evaluated as the mean coordinate of all voxels included. Note that for some of the regions, there are slight inter-individual differences in the exact location of the parcel. This is due to small variations in the normalization of the individual functional images to standard MNI space. Hence, for each region, a somewhat different set of voxels is included in each parcel. This results in a slightly different mean coordinates across all included voxels for each subject. Conversely, for some of the parcels, all participants have the same (full) set of voxels included and, hence, the individual mean coordinates fall onto exactly the same spot. Location of network nodes were visualized using the BrainNet Viewer[14], publicly available for download (http://www.nitrc.org/projects/bnv/). Homotopic regions in the left and the right hemisphere are shown in the same color. L = left hemisphere; R = right hemisphere; A = anterior; P = posterior.

### Supplementary Figure S2

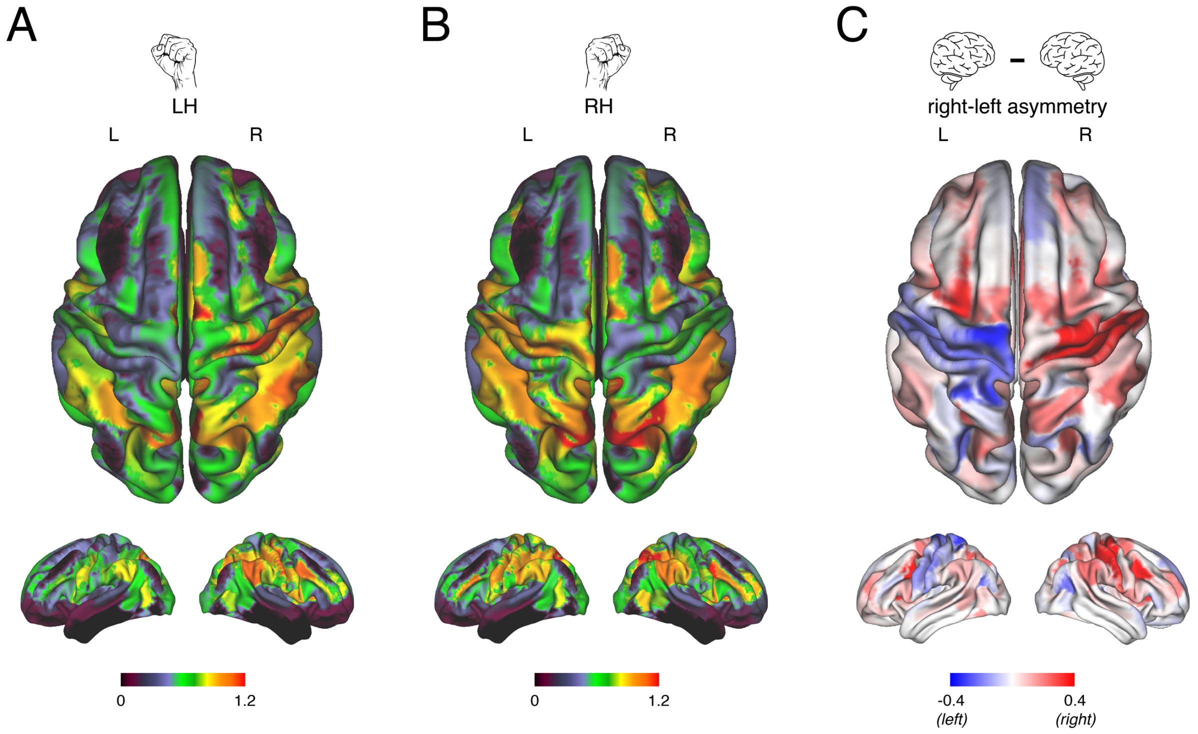


**Supplementary Figure S2:** Graph-theoretical analysis of the whole-brain effective connectivity patterns underlying unilateral hand movements as inferred using rDCM when embedded sparsity constraints were used to prune a fully (all-to-all) connected network. Node strength (i.e., the sum of the weights of all in- and outgoing connections) was evaluated for each parcel of the Human Brainnetome atlas for **(A)** left-hand and **(B)** right-hand fist closings and then graphically projected onto a whole-brain volume. **(C)** Hemispheric asymmetries in node strength were assessed by evaluating the difference in node strength for homotopic parcels in the left and right hemisphere. Positive values indicated higher node strength in the right hemisphere (*red*), whereas negative values indicated higher node strength in the left hemisphere (*blue*). Results for left-hand movements are presented in the right hemisphere, whereas results for right-hand movements are presented in the left hemisphere. Again, this clearly illustrates the mirror symmetry of the motor network in the pre- and postcentral gyrus. Node strength for directed and weighted adjacency matrices was computed using the Brain Connectivity toolbox[15], which is freely available (https://sites.google.com/site/bctnet/). Node strength values for each parcel were visualized using the Human Connectome Workbench, also publicly available (https://www.humanconnectome.org/software/connectome-workbench). L = left hemisphere; R = right hemisphere; LH = left hand; RH = right hand.

**Supplementary Figure S3**

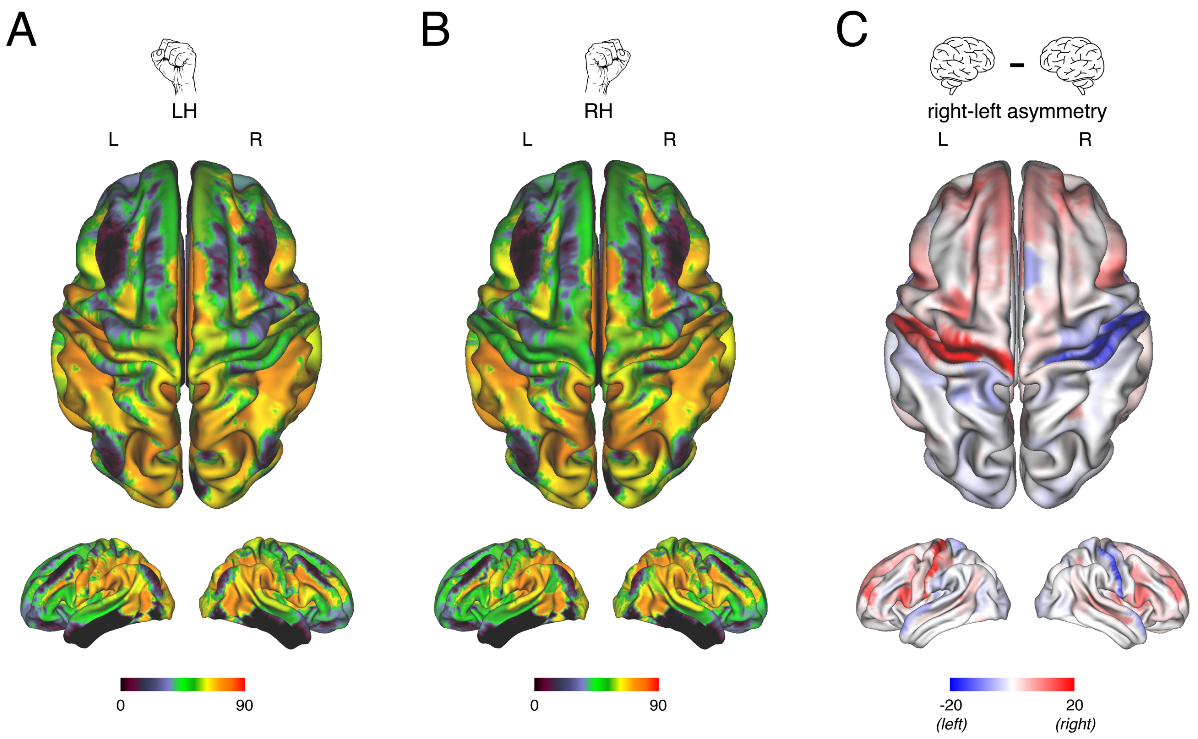


**Supplementary Figure S3:** Graph-theoretical analysis of the functional connectivity patterns underlying unilateral hand movements as assessed using the Pearson correlation coefficient among the same BOLD signal time series as utilized for the rDCM analysis. Node strength (i.e., the sum of the weights of all in- and outgoing connections) was evaluated for each parcel of the Human Brainnetome atlas for **(A)** left-hand and **(B)** right-hand fist closings and then graphically projected onto a whole-brain volume. **(C)** Hemispheric asymmetries in node strength were assessed by evaluating the difference in node strength for homotopic parcels in the left and right hemisphere. Positive values indicated higher node strength in the right hemisphere (*red*), whereas negative values indicated higher node strength in the left hemisphere (*blue*). Results for left-hand movements are presented in the right hemisphere, whereas results for right-hand movements are presented in the left hemisphere. Node strength for undirected and weighted adjacency matrices was computed using the Brain Connectivity toolbox[15], which is freely available (https://sites.google.com/site/bctnet/). Node strength values for each parcel were visualized using the Human Connectome Workbench, also publicly available (https://www.humanconnectome.org/software/connectome-workbench). L = left hemisphere; R = right hemisphere; LH = left hand; RH = right hand.

### Tables

**Supplementary Table S1:** List of brain areas and their abbreviations in the Human Brainnetome atlas[13]. Note that for each region there is a parcel in the left and right hemisphere. Regions from which BOLD signal time series could not be extracted in all participants (see Methods) and which consequently were not included in the rDCM analysis are colored in red.

| Lobe | Gyrus | Abbreviation (left and right hemisphere) | Brainnetome parcel name |
|---|---|---|---|
| **Frontal** | Superior frontal gyrus (SFG) | SFG_L(R)_7_1 | A8m, medial area 8 |
| | | SFG_L(R)_7_2 | A8dl, dorsolateral area 8 |
| | | SFG_L(R)_7_3 | A9l, lateral area 9 |
| | | SFG_L(R)_7_4 | A6dl, dorsolateral area 6 |
| | | SFG_L(R)_7_5 | A6m, medial area 6 |
| | | SFG_L(R)_7_6 | A9m, medial area 9 |
| | | SFG_L(R)_7_7 | A10m, medial area 10 |
| | Middle frontal gyrus (MFG) | MFG_L(R)_7_1 | A9/46d, dorsal area 9/46 |
| | | MFG_L(R)_7_2 | IFJ, inferior frontal junction |
| | | MFG_L(R)_7_3 | A46, area 46 |
| | | MFG_L(R)_7_4 | A9/46v, ventral area 9/46 |
| | | MFG_L(R)_7_5 | A8vl, ventrolateral area 8 |
| | | MFG_L(R)_7_6 | A6vl, ventrolateral area 6 |
| | | MFG_L(R)_7_7 | A10l, lateral area 10 |
| | Inferior frontal gyrus (MFG) | IFG_L(R)_6_1 | A44d, dorsal area 44 |
| | | IFG_L(R)_6_2 | IFS, inferior frontal sulcus |
| | | IFG_L(R)_6_3 | A45c, caudal area 45 |
| | | IFG_L(R)_6_4 | A45r, rostral area 45 |
| | | IFG_L(R)_6_5 | A44op, opercular area 44 |
| | | IFG_L(R)_6_6 | A44v, ventral area 44 |
| | Orbital gyrus (OrG) | OrG_L(R)_6_1 | A14m, media area 14 |
| | | OrG_L(R)_6_2 | A12/47o, orbital area 12/47 |
| | | OrG_L(R)_6_3 | A11l, lateral area 11 |
| | | OrG_L(R)_6_4 | A11m, medial area 11 |

| | | | |
|---|---|---|---|
| | | OrG_L(R)_6_5 | A13, area 13 |
| | | OrG_L(R)_6_6 | A12/47l, lateral area 12/47 |
| | Precentral gyrus (PrG) | PrG_L(R)_6_1 | A4hf, area 4 (head and face region) |
| | | PrG_L(R)_6_2 | A6cdl, caudal dorsolateral area 6 |
| | | PrG_L(R)_6_3 | A4ul, area 4 (upper limb region) |
| | | PrG_L(R)_6_4 | A4t, area 4 (trunk region) |
| | | PrG_L(R)_6_5 | A4tl, area 4 (tongue and larynx region) |
| | | PrG_L(R)_6_6 | A5cvl, causal ventrolateral area 6 |
| | Paracentral lobule (PCL) | PCL_L(R)_2_1 | A1/2/3ll, area 1/2/3 (lower limb region) |
| | | PCL_L(R)_2_2 | A4ll, area 4 (lower limb region) |
| **Temporal** | Superior temporal gyrus (STG) | STG_L(R)_6_1 | A38m, medial area 38 |
| | | STG_L(R)_6_2 | A41/42, area 41/42 |
| | | STG_L(R)_6_3 | TE1.0 and TE1.2 |
| | | STG_L(R)_6_4 | A22c, causal area 22 |
| | | STG_L(R)_6_5 | A38l, lateral area 38 |
| | | STG_L(R)_6_6 | A22r, rostral area 22 |
| | Middle temporal gyrus (MTG) | MTG_L(R)_4_1 | A21c, caudal area 21 |
| | | MTG_L(R)_4_2 | A21r, rostral area 21 |
| | | MTG_L(R)_4_3 | A37dl, dorsolateral area 37 |
| | | MTG_L(R)_4_4 | aSTS, anterior superior temporal sulcus |
| | Inferior temporal gyrus (ITG) | ITG_L(R)_7_1 | A20iv, intermediate ventral area 20 |
| | | ITG_L(R)_7_2 | A37elv, extreme lateroventral area 37 |
| | | ITG_L(R)_7_3 | A20r, rostral area 20 |
| | | ITG_L(R)_7_4 | A20il, intermediate lateral area 20 |
| | | ITG_L(R)_7_5 | A37vl, ventrolateral area 37 |
| | | ITG_L(R)_7_6 | A20cl, caudolateral of area 20 |
| | | ITG_L(R)_7_7 | A20cv, caudoventral of area 20 |
| | Fusiform gyrus (FuG) | FuG_L(R)_3_1 | A20rv, rostroventral area 20 |
| | | FuG_L(R)_3_2 | A37mv, medioventral area 37 |
| | | FuG_L(R)_3_3 | A37lv, lateroventral area 37 |
| | Parahippocampal gyrus (PhG) | PhG_L(R)_6_1 | A35/36r, rostral area 35/36 |
| | | PhG_L(R)_6_2 | A35/36c, caudal area 35/36 |
| | | PhG_L(R)_6_3 | TL, area TL (lateral PPHC, posterior parahippocampal gyrus) |

| | | | |
|---|---|---|---|
| | | PhG_L(R)_6_4 | A28/34, area 28/34 (EC, entorhinal cortex) |
| | | PhG_L(R)_6_5 | TI, area TI (temporal agranular insular cortex) |
| | | PhG_L(R)_6_6 | TH, area TH (medial PPHC) |
| | Posterior superior temporal gyrus (pSTG) | pSTG_L(R)_2_1 | rpSTS, rostroposterior superior temporal sulcus |
| | | pSTG_L(R)_2_2 | cpSTS, caudoposterior superior temporal sulcus |
| **Parietal** | Superior parietal lobule (SPL) | SPL_L(R)_5_1 | A7r, rostral area 7 |
| | | SPL_L(R)_5_2 | A7c, caudal area 7 |
| | | SPL_L(R)_5_3 | A5l, lateral area 5 |
| | | SPL_L(R)_5_4 | A7pc, postcentral area 7 |
| | | SPL_L(R)_5_5 | A7ip, intraparietal area 7 |
| | Inferior parietal lobule (IPL) | IPL_L(R)_6_1 | A39c, causal area 39 (PGp) |
| | | IPL_L(R)_6_2 | A39rd, rostrodorsal area 39 (Hip3) |
| | | IPL_L(R)_6_3 | A40rd, rostrodorsal area 40 (PFt) |
| | | IPL_L(R)_6_4 | A40c, causal area 40 (PFm) |
| | | IPL_L(R)_6_5 | A39rv, rostroventral area 39 (PGa) |
| | | IPL_L(R)_6_6 | A40rv, rostroventral area 40 (PFop) |
| | Precuneus (Pcun) | PCun_L(R)_4_1 | A7m, medial area 7 (PEp) |
| | | PCun_L(R)_4_2 | A5m, medial area 5 (PEm) |
| | | PCun_L(R)_4_3 | dmPOS, dorsomedial parietooccipital sulcus (PEr) |
| | | PCun_L(R)_4_4 | A31, area 31 (Lc1) |
| | Postcentral gyrus (PoG) | PoG_L(R)_4_1 | A1/2/3ulhf, area 1/2/3 (upper limb, head and face region) |
| | | PoG_L(R)_4_2 | A1/2/3tonla, area 1/2/3 (tongue and larynx region) |
| | | PoG_L(R)_4_3 | A2, area 2 |
| | | PoG_L(R)_4_4 | A1/2/3tru, area 1/2/3 (trunk region) |
| **Insular** | Insular gyrus (INS) | INS_L(R)_6_1 | G, hypergranular insula |
| | | INS_L(R)_6_2 | vIa, ventral agranular insula |
| | | INS_L(R)_6_3 | dIa, dorsal agranular insula |
| | | INS_L(R)_6_4 | vId/vIg, ventral dysgranular and granular insula |
| | | INS_L(R)_6_5 | dIg, dorsal granular insula |
| | | INS_L(R)_6_6 | dId, dorsal dysgranular insula |

| | | | |
|---|---|---|---|
| **Limbic** | Cingulate gyrus (CG) | CG_L(R)_7_1 | A23d, dorsal area 23 |
| | | CG_L(R)_7_2 | A24rv, rostroventral area 24 |
| | | CG_L(R)_7_3 | A32p, pregenual area 32 |
| | | CG_L(R)_7_4 | A23v, ventral area 23 |
| | | CG_L(R)_7_5 | A24cd, caudodorsal area 24 |
| | | CG_L(R)_7_6 | A23c, caudal area 23 |
| | | CG_L(R)_7_7 | A32sg, subgenual area 32 |
| **Occipital** | Medioventral occipital cortex (MVOoC) | MVOoC_L(R)_5_1 | cLinG, caudal lingual gyrus |
| | | MVOoC_L(R)_5_2 | rCunG, rostral cuneus gyrus |
| | | MVOoC_L(R)_5_3 | cCunG, caudal cuneus gyrus |
| | | MVOoC_L(R)_5_4 | rLinG, rostral lingual gyrus |
| | | MVOoC_L(R)_5_5 | vmPOS, ventromedial parietooccipital sulcus |
| | Lateral occipital cortex (LOoC) | LOoC_L(R)_4_1 | MOccG, middle occipital gyrus |
| | | LOoC_L(R)_4_2 | V5/MT+, area V5/MT+ |
| | | LOoC_L(R)_4_3 | OPC, occipital polar cortex |
| | | LOoC_L(R)_4_4 | iOccG, inferior occipital gyrus |
| | | LOoC_L(R)_2_1 | msOccG, medial superior occipital gyrus |
| | | LOoC_L(R)_2_2 | lsOccG, lateral superior occipical gyrus |
| **Subcortical** | Amygdala (Amyg) | Amyg_L(R)_2_1 | mAmyg, medial amygdala |
| | | Amyg_L(R)_2_2 | lAmyg, lateral amygdala |
| | Hippocampus (Hipp) | Hipp_L(R)_2_1 | rHipp, rostral hippocampus |
| | | Hipp_L(R)_2_2 | cHipp, caudal hippocampus |
| | Basal ganglia (BG) | BG_L(R)_6_1 | vCa, ventral caudate |
| | | BG_L(R)_6_2 | GP, globus pallidus |
| | | BG_L(R)_6_3 | NAC, nucleus accumbens |
| | | BG_L(R)_6_4 | vmPu, ventromedial putamen |
| | | BG_L(R)_6_5 | dCa, dorsal caudate |
| | | BG_L(R)_6_6 | dlPu, dorsolaterael putamen |
| | Thalamus (Tha) | Tha_L(R)_8_1 | mPFtha, medial pre-frontal thalamus |
| | | Tha_L(R)_8_2 | mPMtha, pre-motor thalamus |
| | | Tha_L(R)_8_3 | Stha, sensory thalamus |

| | |
|---|---|
| Tha_L(R)_8_4 | rTtha, rostral temporal thalamus |
| Tha_L(R)_8_5 | PPtha, posterior parietal thalamus |
| Tha_L(R)_8_6 | Otha, occipital thalamus |
| Tha_L(R)_8_7 | cTtha, causal temporal thalamus |
| Tha_L(R)_8_8 | lPFtha, lateral pre-frontal thalamus |

**Supplementary Table S2:** Peak coordinates and T-scores of BOLD activations related to visually synchronized left- and right-hand movements. Anatomical labels have been identified using the Anatomy toolbox extension[16]. Listed are regions that were significantly activated at a voxel-level threshold $p<0.05$ (FWE-corrected) and for which the cluster size exceeded 10 voxels.

| Cortical region | Hemisphere | MNI coordinates | | | t score |
|---|---|---|---|---|---|
| | | x | y | z | |
| *Left-hand movements* | | | | | |
| Precentral gyrus (**M1**) | R | 48 | -20 | 59 | 14.83 |
| Postcentral gyrus | R | 40 | -24 | 52 | 14.44 |
| Cerebellum | L | -16 | -50 | -22 | 13.39 |
| Putamen | R | 34 | -4 | -4 | 12.53 |
| Suppl. motor area (**SMA**) | L | -6 | -4 | 56 | 11.27 |
| Precentral gyrus (**M1**) | L | -34 | -13 | 62 | 10.82 |
| Precentral (**PMC**) | R | 52 | 6 | 44 | 9.99 |
| Middle occipital (**V5**) | R | 30 | -88 | -2 | 9.96 |
| Suppl. motor area (**SMA**) | R | 4 | 2 | 62 | 9.75 |
| Precentral (**PMC**) | L | -51 | 0 | 48 | 9.81 |
| Thalamus | R | 18 | -19 | 2 | 9.69 |
| Cerebellum | R | 30 | -49 | -25 | 9.09 |
| Pars oppercularis | L | -54 | 6 | 18 | 8.65 |
| Rolandic operculum | R | 50 | -22 | 23 | 8.63 |
| Postcentral gyrus | L | -57 | -20 | 32 | 8.03 |
| Thalamus | L | -9 | -16 | 2 | 7.80 |
| Posterior medial frontal | L | -6 | 5 | 48 | 7.62 |
| *Right-hand movements* | | | | | |
| Postcentral | L | -36 | -26 | 56 | 23.35 |
| Precentral gyrus (**M1**) | L | -39 | -22 | 53 | 22.32 |
| Cerebellum | R | 22 | -52 | -20 | 18.17 |
| Suppl. motor area (**SMA**) | L | -6 | -10 | 56 | 15.97 |
| Thalamus | L | -14 | -20 | 4 | 12.60 |
| Suppl. motor area (**SMA**) | R | 3 | 2 | 62 | 11.28 |
| Middle occipital (**V5**) | R | 30 | -84 | 0 | 11.26 |
| Precentral (**PMC**) | R | 58 | 12 | 34 | 9.98 |
| Rolandic operculum | L | -45 | -26 | 22 | 9.90 |
| Precentral (**PMC**) | L | -50 | -1 | 38 | 9.61 |
| Middle occipital (**V5**) | L | -42 | -73 | -10 | 9.25 |
| Inferior parietal lobe | R | 30 | -49 | 47 | 8.83 |
| Cerebellum | L | -27 | -54 | -25 | 8.45 |
| Rolandic operculum | R | 51 | 6 | 12 | 8.18 |
| Fusiform gyrus | R | 36 | -58 | -10 | 8.09 |
| Inferior parietal lobe | L | -27 | -49 | 47 | 8.07 |
| Superior temporal gyrus | L | -54 | -36 | 22 | 7.88 |
| Putamen | L | -27 | -2 | 11 | 7.69 |
